# Individual connectome fingerprints reveal early stabilization and long-term circuit remodeling after stroke

**DOI:** 10.64898/2026.03.27.714711

**Authors:** Andrea Santoro, Alessandro Lucatelli, Fabienne Windel, Beatrice Lugli, Maria Giulia Preti, Lisa Fleury, Flavia Petruso, Elena Beanato, Dimitri Van De Ville, Friedhelm Christoph Hummel, Enrico Amico

## Abstract

Stroke is one of the leading causes of global disability, yet the principles governing how focal brain injury disrupts large-scale neural connectivity over time remain poorly understood. Here, we leverage a longitudinal multimodal dataset to track the evolution of individual-specific connectivity patterns, or ‘brain fingerprints’, over the first year after stroke. Despite a persistent shift from healthy architecture, we demonstrate that each patient’s unique functional connectome fingerprint is remarkably resilient and stabilizes within three weeks. This early global stabilization masks a protracted system-specific reorganization of brain circuits, which is characterized by an initial increase in connectivity within sensory and attention systems, followed by a decline across higher-level association networks. A joint structure–function embedding further shows that recovery involves a gradual shift toward the normative healthy range, driven primarily by functional reconfiguration atop a stable structural lesion. Crucially, a multivariate prediction model reveals that early functional signatures selectively forecast long-term impairment in language, executive function, and attention. Together, our results define the post-stroke brain as a shifting but constrained dynamical system, identifying early-stabilized brain patterns as biomarkers for individual recovery profiles and targets for personalized neurorehabilitation.

---

Advancements in network neuroscience have revolutionized our understanding of the brain [1, 2], shifting from the concept of isolated regions to recognizing it as a highly interconnected system [3, 4]. The availability of large-scale neuroimaging datasets [5, 6] has enabled detailed mapping of functional and structural connections, providing invaluable benchmarks for exploring neural connectivity [7, 8]. This network-based perspective has yielded profound insights into how different brain regions interact to support brain function and how their disruption contributes to neurological disorders [9–12].

A notable development in this field is the concept of brain fingerprinting, which posits that an individual’s unique pattern of brain connectivity can serve as a distinctive identifier, much like a fingerprint [13–16]. Early studies demonstrated that functional connectomes derived from resting-state functional magnetic resonance imaging (fMRI) are stable over time and can reliably distinguish individuals within a population [17]. Beyond fMRI, brain fingerprinting has been observed across modalities — including EEG, MEG and fNIRS — supporting the notion that stable, person-specific features are a general property of brain organization rather than an artifact of a single measurement technique [18–23]. These individualized profiles have also been linked to individual differences in cognitive abilities and behavioral traits [24–27], underscoring the potential of brain finger-printing as a biomarker for personalized neuroscience. Yet, despite increasing applications in neuropsychiatric and neurodegenerative contexts [28–31], the extent to which fingerprints persist, reorganize, or break down after acute focal brain injury remains largely unknown.

Stroke offers a compelling clinical case. It is a leading cause of disability worldwide [32–34], and is increasingly understood as a network disorder: a focal lesion can trigger widespread changes in both structural and functional connectivity through direct disconnection and remote physiological effects [35–43]. Mechanisms such as diaschisis [44], through which localized brain injury causes dysfunction in remote regions due to disrupted neural pathways, illustrate how stroke-induced damage extends beyond the lesion, affecting widespread neural pathways. This extensive network disruption impairs communication between brain regions [45–47], altering local and global network properties [48–50] and impacting both cognitive and motor functions [51–56].

The potential for functional recovery following stroke varies considerably among patients [57, 58], with fewer than 15% achieving complete motor restoration [59] and pooled analyses reporting post-stroke cognitive impairment in approximately 38% of individuals [60]. This variability has been linked to lesion size, lesion location, damage to critical white matter tracts, and individual neuroplastic responses [42, 61–63]. Neural reorganization after stroke is a dynamic, time-sensitive process involving both the ipsilesional and contralesional hemispheres. While an early increase of contralesional connectivity may initially serve a compensatory role, prolonged hyperactivity in the contralesional hemisphere has been associated with poorer long-term outcomes, possibly due to maladaptive plasticity interfering with recovery in the affected hemisphere [64]. Conversely, strengthening ipsilesional connectivity and promoting reorganization within the damaged hemisphere are often correlated with better functional recovery [65]. These opposing neural changes underscore the importance of precisely characterizing connectivity patterns that facilitate (or hinder) long-term rehabilitation. Despite these insights, most studies to date rely on group-level analyses or cross-sectional designs, which fail to capture individual-level trajectories and the evolving nature of recovery [49, 66, 67]. Additionally, it remains unclear how functional connectivity patterns evolve from the acute phase to chronic stages, and how these changes relate to clinical outcomes [52, 68–70]. These gaps highlight the need for longitudinal investigations that can more accurately map the interplay between network reorganization and functional gains.

Here, we address these open questions using a longitudinal multimodal dataset [71], tracking stroke patients across four distinct stages during the first year after the ictal event. We combine complementary analyses to conceptualize this recovery as an individualized, time-resolved network process. We first quantify the resilience of patient-specific functional “fingerprints” using identifiability tools [17], contextualizing these stable signatures against a largely stationary structural disconnection backbone and evolving functional hyper- and hypo-connectivity. We then embed patients in a joint structure–function state space, framing recovery as a shift toward healthy variability under strict anatomical constraints. Finally, using multivariate brain–behavior association and ridge-based predictive modeling, we test whether early functional signatures forecast long-term clinical recovery across multiple domains. Together, this framework moves beyond group averages to connect stable individual identity with selective circuit remodeling, providing a robust route toward individualized biomarkers for prognosis and targeted neurorehabilitation.

## RESULTS

To investigate the principles governing individual brain network reorganization after a focal injury, we leveraged the TiMeS cohort — a longitudinal, observational study of stroke recovery characterized by high-density multimodal imaging, including resting-state fMRI and diffusion-weighted imaging, as well as a detailed clinical assessment [71]. We analyzed *N* = 64 patients across four critical stages of recovery: acute (*T*_1_, ∼ 1 week), early sub-acute (*T*_2_, ∼ 3 weeks), late sub-acute (*T*_3_, ∼ 3 months), and chronic (*T*_4_, ∼ 12 months). Lesion anatomy was heterogeneous but frequently involved subcortical structures (Fig. 1**a**), providing a stringent test of how disruption of cortico-subcortical pathways reshapes large-scale functional organization over time. To maximize statistical power, all available data were included for each analysis unless otherwise specified (see Methods).

**FIG. 1.**
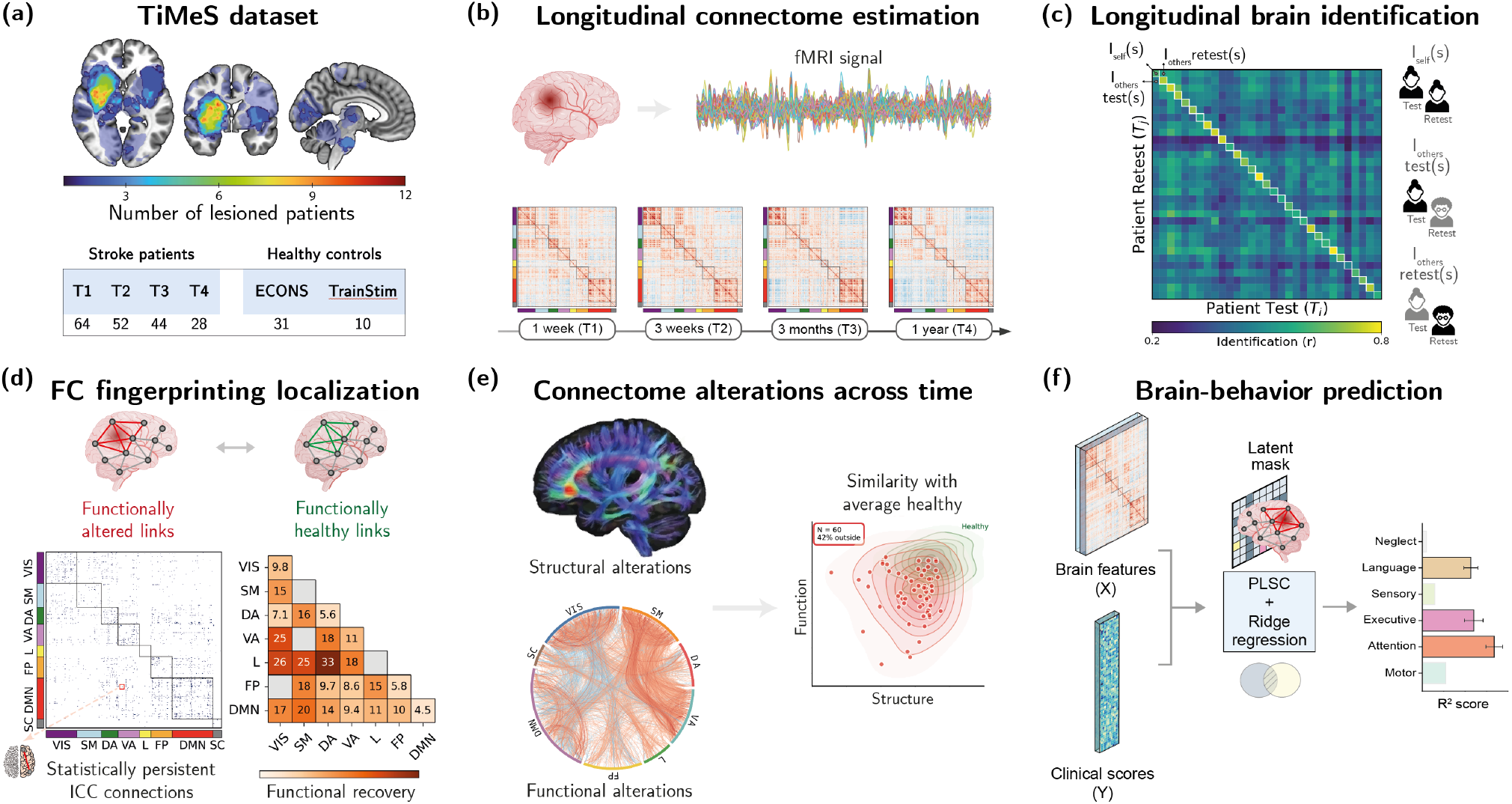
Mapping connectome disruption and individual stability after stroke. **(a)** TiMeS cohort [71], showing the lesion overlap map for the stroke cohort and sample sizes across four post-stroke assessments (T1: ∼ 1 week; T2: ∼ 3 weeks; T3: ∼ 3 months; T4: ∼ 1 year), together with healthy control groups used for benchmark (ECONS and TrainStim). **(b)** Resting-state fMRI BOLD time series were used to derive whole-brain functional connectivity (FC) matrices at each time point for each participant. **(c)** Longitudinal brain identification based on a test–retest similarity matrix obtained by correlating each patient’s connectivity pattern at test (*T*_*i*_) and retest (*T*_*j*_) sessions; high diagonal values (*I*_self_ ) indicate reliable within-subject connectivity “fingerprints” relative to between-subject similarity (*I*_others_). **(d)** Spatial specificity analysis of the functional connectome fingerprint, in which statistically persistent connections (intraclass correlation, ICC) are classified as functionally altered or functionally preserved and summarized across canonical large-scale networks to map patterns of recovery. **(e)** Structural disconnection and functional deviations relative to healthy controls are quantified longitudinally and summarized in a joint structure–function similarity space, capturing patients’ recovery trajectories. **(f)** Brain–behavior associations were assessed using multivariate Partial Least Squares Correlation (PLSC) to identify connectivity patterns covarying with clinical scores; the resulting latent connectivity mask was then used to constrain ridge regression models for out-of-sample prediction of longitudinal outcomes across behavioral domains.

Recognizing that group-level averages often conceal the idiosyncratic nature of recovery, we developed a multi-stage analytical framework to track the functional reconfiguration of each patient’s connectome over time (Fig. 1**b–f** ). For each session, whole-brain functional connectivity (FC) was estimated from resting-state BOLD time series using a parcellation of 377 regions, comprising 360 cortical areas from the Glasser atlas [72] and 17 subcortical regions (see Methods). Connectivity patterns were benchmarked against a normative reference derived from two healthy control cohorts acquired with matched scanning protocols and preprocessing (ECONS and TrainStim, for a total of *n* = 41 subjects; see Methods).

To quantify the preservation of individual-specific topography despite injury, we employed a brain fingerprinting approach [17]. For each session pairing (*T*_*i*_, *T*_*j*_), we constructed an identifiability matrix and derived within-subject similarity (*I*_self_ ) and between-subject similarity (*I*_others_) to assess whether patients remained more similar to their own past or future selves than to other individuals (Fig. 1**c**). Furthermore, we localized stability and remodeling by estimating link-wise reliability with intraclass correlation (ICC) [73, 74], distinguishing statistically persistent connections from those that were selectively dynamic across recovery and summarizing these effects across canonical functional systems (Fig.1**d**).

We then characterized structural damage and functional dysfunction at complementary scales, from individual links to canonical network blocks, and summarized each patient’s longitudinal evolution in a joint structure– function space (Fig. 1**e**). In this embedding, patients were positioned according to their structural and functional similarity to the healthy reference, providing a compact description of how whole-brain states shift relative to the range of healthy variability and motivating a dynamical-systems view of recovery as transitions within a constrained landscape. Finally, to establish the clinical relevance of these network shifts, we coupled connectome reconfiguration to multidomain behavioral profiles using a two-stage framework (Fig. 1**f** ). Partial Least Squares Correlation (PLSC) [75] identified latent brain–behavior components at a given time point, and the resulting connectivity signature was carried forward to constrain ridge regression models for out-of-sample prediction of followup clinical scores while avoiding data-leakage from feature selection [76, 77] (see Methods for details).

### The individual connectome fingerprint is resilient and consolidates early after stroke

We first investigated whether focal injury induces a return toward canonical healthy organization or a persistence in a pathological state. Using *I*_clinical_, we quantified the similarity of each patient’s FC to the healthy reference cohort (ECONS) and benchmarked these values against the baseline similarity observed between two independent healthy groups (TrainStim vs. ECONS; Fig. 2**a**). Across the first year post-stroke, patients exhibited a sustained and significant shift away from the healthy normative range. This divergence was robust in the acute (*T*_1_), early sub-acute (*T*_2_), and chronic stages (*T*_4_; *p <* 0.01, FDR-corrected Mann-Whitney U test), while the *T*_3_ interval exhibited a marginal trend (*p* = 0.053).

**FIG. 2.**
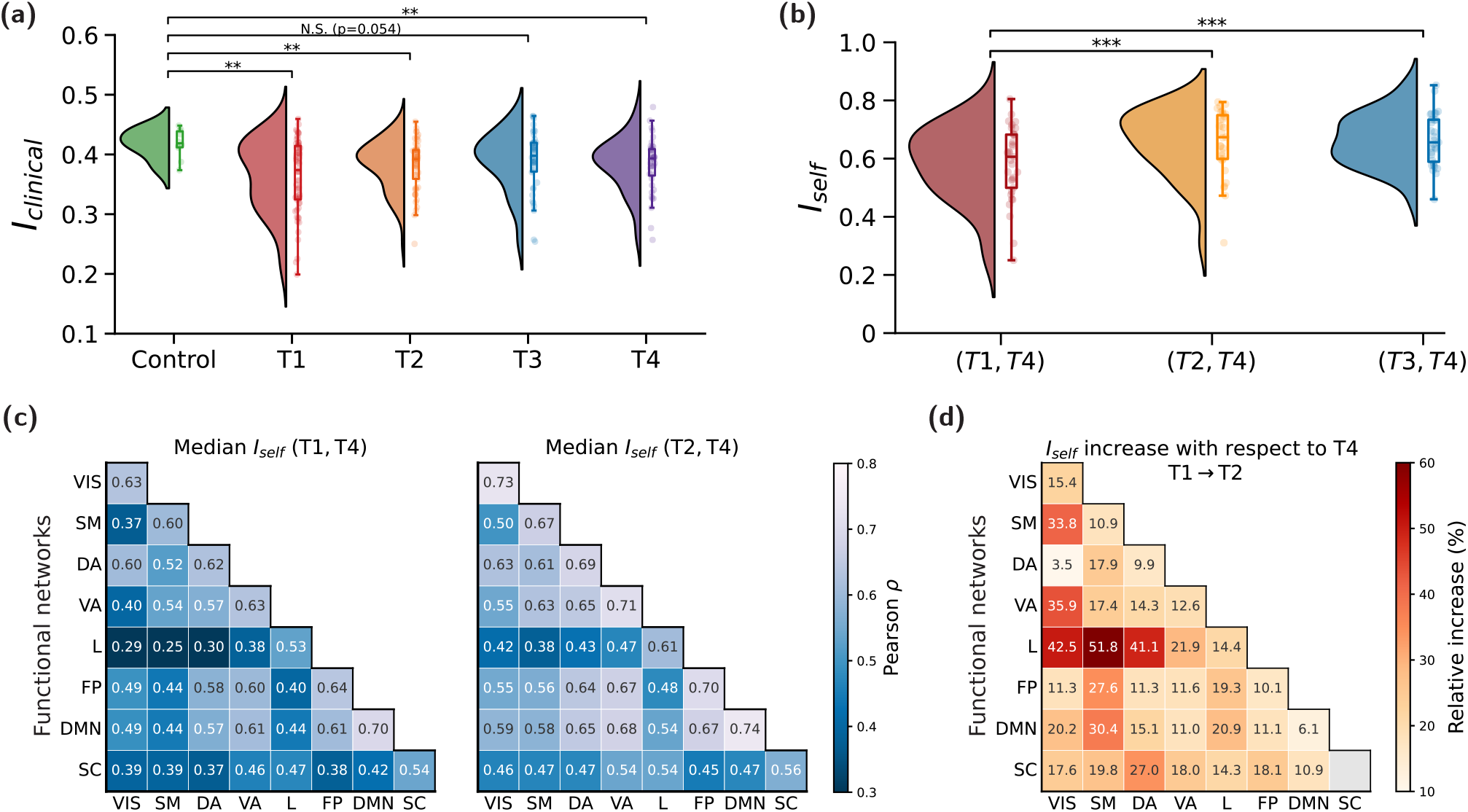
Longitudinal evolution of functional connectome identification following stroke. **(a)** Distribution of connectome similarity scores *I*_clinical_ for healthy controls (TrainStim vs. ECONS) and stroke patients (relative to ECONS) across four time points. Stroke scores are significantly lower than healthy controls at *T*_1_, *T*_2_, and *T*_4_ ( ∗∗*p <* 0.01, FDR-corrected Mann-Whitney U test), with a marginal trend at *T*_3_ (*p* = 0.053). **(b)** Distribution of self-identifiability scores (*I*_self_ ), quantifying patient connectome similarity between earlier time points and the chronic outcome (T4). Significantly lower *I*_self_ at *T*_1_ compared to *T*_2_ indicates rapid functional reorganization of the individual fingerprint within the first three weeks post-stroke ( ∗∗∗*p <* 0.001, LMM). *I*_self_ at *T*_2_ shows no significant difference from *T*_3_ (*p*_FDR_ = 0.054). **(c)** Median *I*_self_ within and between seven canonical networks and subcortical structures (VIS, visual; SM, somatomotor; DA, dorsal attention; VA, ventral attention; L, limbic; FP, frontoparietal; DMN, default mode) and subcortical structures (SC). Comparing (T1,T4) and (T2,T4) reveals how specific networks consolidate their functional distinctiveness over time. **(d)** Significant relative percentage increase in *I*_self_ from acute to sub-acute stages (*T*_1_ → *T*_2_), referenced to the chronic baseline (*T*_4_). Warmer colors denote networks undergoing rapid functional reconfiguration, notably the ventral attention (VA), limbic (L), and somatomotor (SM) systems (*p <* 0.05, LMM).

Despite this sustained global shift, FC patterns of stroke patients remained strongly identifiable. *I*_self_ was consistently higher than between-subject similarity, demonstrating that each patient preserved an idiosyncratic connectome signature that generalizes across months of recovery (Fig. 2**b**). This individual fingerprint was highly robust even in the acute phase, with identification rates exceeding 75% between *T*_1_ and chronic stages (i.e., *T*_3_, *T*_4_; see Supplementary Fig. S1). This indicates that focal injury perturbs the connectome without erasing its patient-specific organization.

The longitudinal trajectory of identification suggested rapid consolidation rather than prolonged instability. Although the acute phase (*T*_1_) showed the lowest self-similarity relative to the chronic stage, the fingerprint sharpened significantly by the three-week mark (*T*_2_), where identification rates rose to 80%. To account for unbalanced longitudinal sampling and repeated measures, we tested these effects using a linear mixed model (LMM). The LMM confirmed an early stabilization of the fingerprint, confirming that individual signatures stabilize rapidly; *I*_self_ at *T*_2_ showed no significant difference from *T*_3_ when compared to the one-year endpoint of the study (Fig. 2b; LMM, *p*_FDR_ *<* 0.001 for the (*T*_2_, *T*_4_) vs. (*T*_3_, *T*_4_) comparison). These results support a rapid transition toward a stable, individual-specific “recovery brain fingerprint” within the first month, even though the functional connectome remains globally shifted away from healthy benchmark.

This consolidation was not spatially uniform across the connectome. When examining the *I*_self_ distributions on pairs of canonical resting-state networks [78], association networks showed the strongest longitudinal stability (Fig. 2**c**). In particular, the default mode and frontoparietal networks maintained high within-network self-similarity across recovery, consistent with a role as stable anchors of individual functional identity despite prevalent subcortical injury. Converging evidence comes from our edge-wise reliability analysis; using an intraclass-correlation (ICC) approach to identify fingerprint-defining connections, edges within and involving frontoparietal and default mode systems were among the most reliably expressed across sessions (Supplementary Fig. S2), reinforcing their role as “identity-stabilizing” components. By contrast, network blocks involving subcortical and somatomotor-related systems showed lower reliability, consistent with greater susceptibility of cortico-subcortical loops to lesion-driven disruption and compensatory reconfiguration.

Critically, early improvements in identifiability (*I*_self_ ) were localized to specific subsets of functional networks rather than occurring globally (Fig. 2**d**). This suggests that fingerprint consolidation relies on the targeted re-stabilization of specific circuits rather than uniform brain-wide normalization. This rapid shift toward a stable regime by *T*_2_ was highly consistent across different time-point comparisons, confirming that networks are unstable at *T*_1_ but remain stable from *T*_2_ onward (Supplementary Fig. S3). These localized effects were fully preserved in a sensitivity analysis restricted to patients with complete four-timepoint data (Supplementary Fig. S4).”

### Decoupling of functional reconfiguration from structural lesion

The rapid consolidation of whole-brain fingerprints by the early sub-acute stage (*T*_2_) raises a key question: can stable individual-level identity coexist with ongoing remodeling of specific connections? We first quantified the structural impact of stroke by estimating each patient’s disconnectome from diffusion-weighted imaging, capturing the set of white-matter pathways disrupted by the lesion. To minimize longitudinal confounds, we focused on the subset of patients with complete multi-modal data across time. We then summarized the proportion of structurally lesioned edges within and between canonical Yeo networks (Fig. 3**a**, top; orange scale). Remarkably, the overarching topography of structural disconnection remained largely stable over the first year. While this macro-scale pattern was preserved, it is important to note that structural connectivity can undergo secondary longitudinal alterations, such as Wallerian degeneration, particularly in initially damaged networks or following larger lesions [42]. Nevertheless, this conclusion held across alternative lesion-tracking approaches (Supplementary Fig. S5 for comparisons), including lesion-only disconnection estimates and BCB toolkit-based reconstructions [79]. Together, these analyses delineate a stable structural “backbone” that constrains the space of possible functional configurations, yet does not by itself explain or predict the trajectory of functional reorganization. Rather, functional connectivity appears to decouple from the structural lesion backbone over time.

**FIG. 3.**
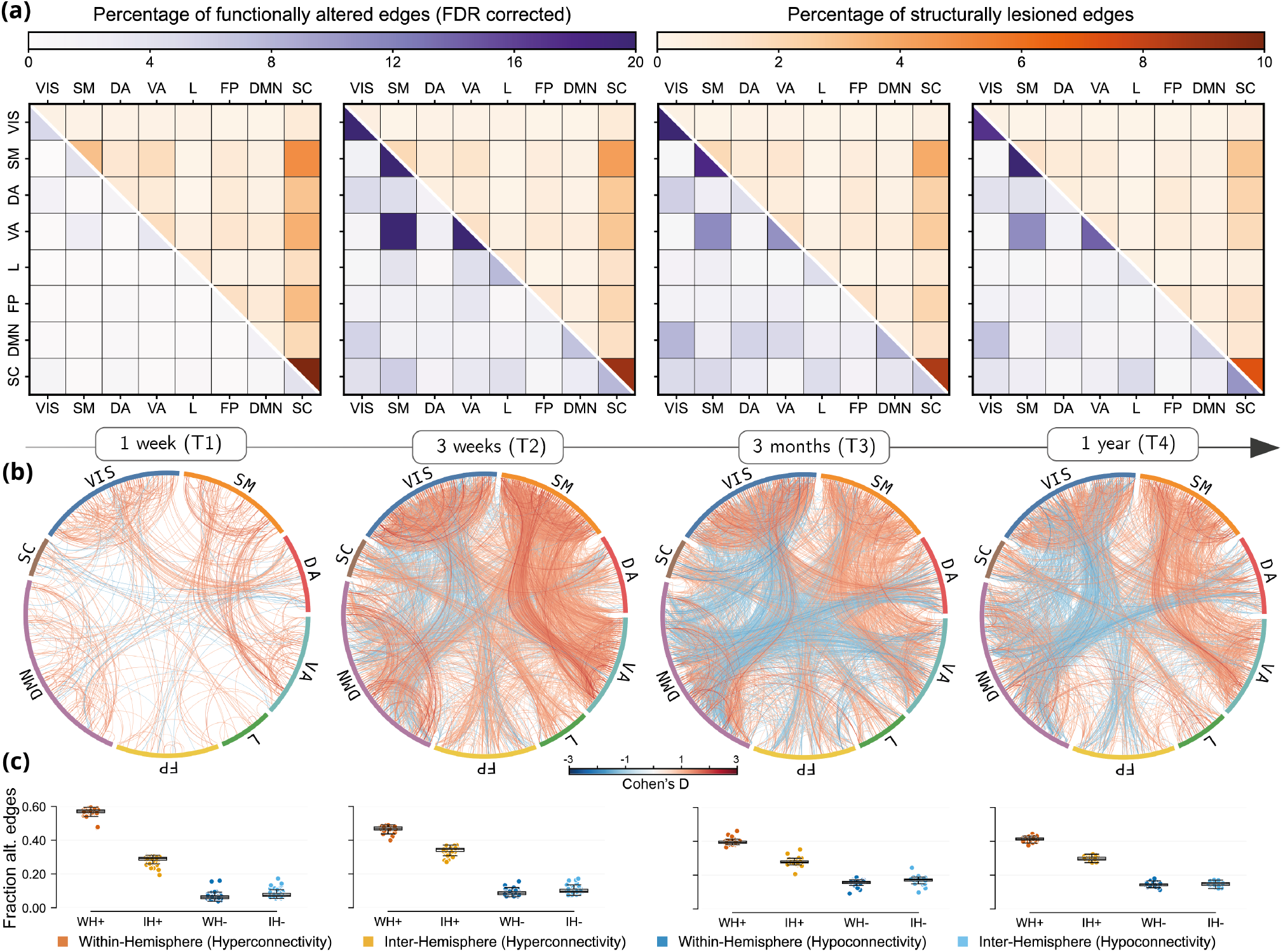
Longitudinal functional reconfiguration decouples from structural damage following stroke. **(a)** Group-level matrices displaying the percentage of DWI-derived structurally lesioned edges (upper triangle) and functionally altered edges (lower triangle; FDR-corrected Mann-Whitney U test, *q <* 0.05) relative to healthy controls across four time points (*T*_1_–*T*_4_). Structural disconnection patterns remain relatively static, whereas functional alterations evolve dynamically, revealing a spatial dissociation between the fixed structural injury and time-varying functional reorganization throughout the first year. **(b)** Circular plots depict positive (hyperconnectivity, red) and negative (hypoconnectivity, blue) functional alterations at T1–T4, as quantified by Cohen’s d effect size. **(c)** Box plots summarize the fraction of altered edges that are within-hemisphere (WH) or inter-hemisphere (IH), further subdivided into hyperconnectivity (*WH*+; *IH*+) and hypoconnectivity (*WH*− ; *IH*− ). Early stages (*T*_1_–*T*_2_) are dominated by within-hemisphere hyperconnectivity, whereas the late sub-acute/chronic phase (*T*_3_–*T*_4_) shows a relative increase in inter-hemispheric alterations, consistent with a shift from local to cross-hemispheric plasticity. Data are shown for the longitudinal subset (*n* = 22). Network abbreviations: VIS, visual; SM, somatomotor; DA, dorsal attention; VA, ventral attention; L, limbic; FP, frontoparietal; DMN, default mode; SC, subcortical.

Indeed, in contrast to this structurally stable disconnection pattern, the functional connectome exhibited a sustained and system-specific trajectory of changes (Fig. 3**a**, bottom; purple scale). To map these alterations without imposing *a priori* constraints, we performed a large-scale univariate analysis across all functional connections (*K* = 70 876), comparing the distribution of each edge in patients versus healthy controls (ECONS) at each time point (Mann–Whitney U test; FDR-corrected *P <* 0.05). Aggregating significant effects within and between Yeo networks revealed a clear decoupling across scales. Although *I*_self_ indicates that global identity stabilizes by three weeks, the set of altered functional links continues to evolve across months, indicating ongoing circuit-level remodeling well into the chronic phase.

Statistical significance identifies which edges differ from healthy norms, but not whether they are strengthened or weakened. We therefore quantified effect sizes (Cohen’s *d*) [80] for all significantly altered links, interpreting positive values (*d >* 0) as hyper-connectivity and negative values (*d <* 0) as hypo-connectivity. Network-level patterns were visualized with circular connectograms and hemispheric summaries (Fig. 3**b–c**). This decomposition revealed a marked temporal asymmetry. Hyper-connectivity was most prominent early after stroke, spanning both intra- and inter-hemispheric connections and disproportionately involving sensory and attention-related systems, consistent with transient compensatory coupling. In contrast, hypo-connectivity accumulated more gradually and became increasingly prevalent from the acute through late sub-acute stages, particularly for links involving higher-order association systems.

These sign-dependent effects were especially evident in the network-resolved profiles. From three months onward (*T*_3_), hyper-connectivity was most pronounced within somatomotor circuitry and in somatomotor–ventral attention (VA) interactions, suggesting strengthened coupling between action-related and salience/attention pathways during late sub-acute recovery. In parallel, widespread hypo-connectivity emerged from FP, DMN, and SC systems, consistent with persistent disruption of cortico-subcortical integration and reduced long-range coupling within association architecture. At the hemispheric level, the fraction of hypo-connected edges increased from *T*_1_ to *T*_3_ (Fig. 3**c**), indicating that negative deviations from the healthy baseline can accumulate even as the patient-specific fingerprint consolidates.

Together, these results establish a dissociation between a largely time-invariant structural injury backbone and a time-varying functional phenotype. The post-stroke connectome rapidly converges to a stable, individually specific configuration at the whole-brain level, yet continues to reorganize through selective, network-specific remodeling.

### Temporal asymmetry of hyper- and hypo-connectivity phenotypes

To quantify the sign-dependent effects more directly, we summarized the *fraction* of significantly altered edges within and between canonical networks, separately for positive and negative deviations from healthy controls, and further stratified them by hemispheric topology (left–left, right–right, and inter-hemispheric blocks; Fig. 4). This representation provides a compact view of how abnormal coupling evolves across recovery while retaining information about network identity and laterality.

**FIG. 4.**
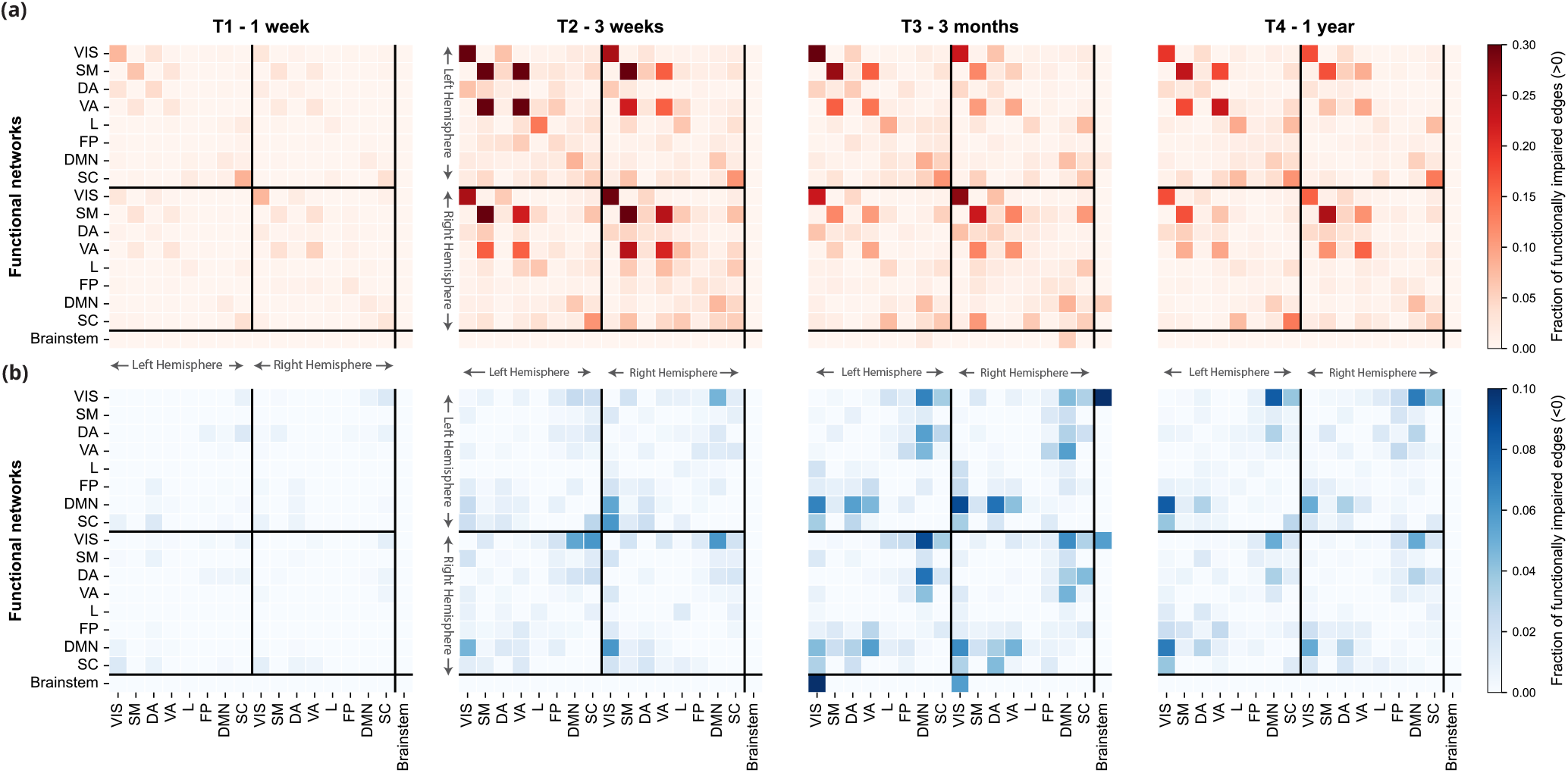
Longitudinal distribution of positive and negative functional network alterations. Matrices report the network-level fraction of functional edges significantly altered in stroke patients relative to healthy controls (ECONS) across four stages (T1: 1 week; T2: 3 weeks; T3: 3 months; T4: 1 year). Significance was determined via edge-wise Mann-Whitney U tests (70 876 comparisons, FDR-corrected at *q <* 0.05), and aggregated by network block. **(a)** Fraction of positive alterations (hyper-connectivity), highlighting a left-hemispheric intra-hemispheric peak at T2, primarily involving VA, L, and FP systems. **(b)** Fraction of negative alterations (hypo-connectivity), which peaks at T3 and persists into the chronic phase, with prominent involvement of VIS and SM networks. Black lines delineate left-hemisphere, right-hemisphere, and inter-hemispheric blocks. Abbreviations: VIS, visual; SM, somatomotor; DA, dorsal attention; VA, ventral attention; L, limbic; FP, frontoparietal; DMN, default mode; SC, subcortical.

Hyper-connectivity followed an early and transient course (Fig. 4**a**). At one week post stroke (*T*_1_), only a small fraction of edges showed increased coupling relative to controls. By three weeks (*T*_2_), hyper-connectivity rose sharply and organized into coherent network blocks, dominated by within-hemisphere effects with additional inter-hemispheric involvement. These increases were most evident among sensory and attention-related systems, consistent with a short-lived phase of compensatory strengthening of functional coupling after the acute insult. By *T*_3_ and *T*_4_, the overall extent of hyper-connectivity decreased, indicating that the early increase does not persist but gives way to a more selective pattern as recovery progresses.

Hypo-connectivity displayed a delayed and more sustained trajectory (Fig. 4**b**). Negative deviations from healthy controls were modest at *T*_1_ but became more prevalent by *T*_2_ and reached their broadest expression at three months (*T*_3_), spanning a wider set of network interactions and involving both intra- and inter-hemispheric edges. These effects were not confined to a single system. Instead, they extended across higher-order association networks and prominently involved subcortical interactions, consistent with persistent disruption of cortico-subcortical integration and long-range coupling. Although the overall burden of hypo-connectivity partially receded by one year (*T*_4_), substantial negative deviations remained, indicating that chronic-stage functional architecture stays measurably altered even after consolidation of global fingerprinting.

These hemispherically resolved trajectories refine the picture emerging from Fig. 3. Post-stroke reorganization is marked by an early increase in hyper-connectivity that peaks in the early sub-acute stage, followed by a broader rise in hypo-connectivity that culminates months after the event. This temporal asymmetry supports a model in which the connectome stabilizes rapidly at the level of global identity, while specific network interactions continue to be reshaped through sequential, partly opposing phases of functional coupling during recovery.

### Recovery as a longitudinal migration toward a healthy structure–function manifold

The results above point to a dissociation across scales. Whole-brain identifiability stabilizes early, yet specific links and network interactions continue to remodel over months. We therefore asked whether these high-dimensional changes can be summarized as a coordinated shift in joint structure–function organization (Fig. 5). This view frames recovery as the evolution of a dynamical system in which large-scale functional activity is constrained by an attractor-like landscape shaped by the structural connectome [81–83]. We embedded each patient at each time point in a two-dimensional state space defined by similarity of their structural and functional connectome to the mean healthy template (ECONS), and benchmarked patient positions against the distribution of healthy variability.

**FIG. 5.**
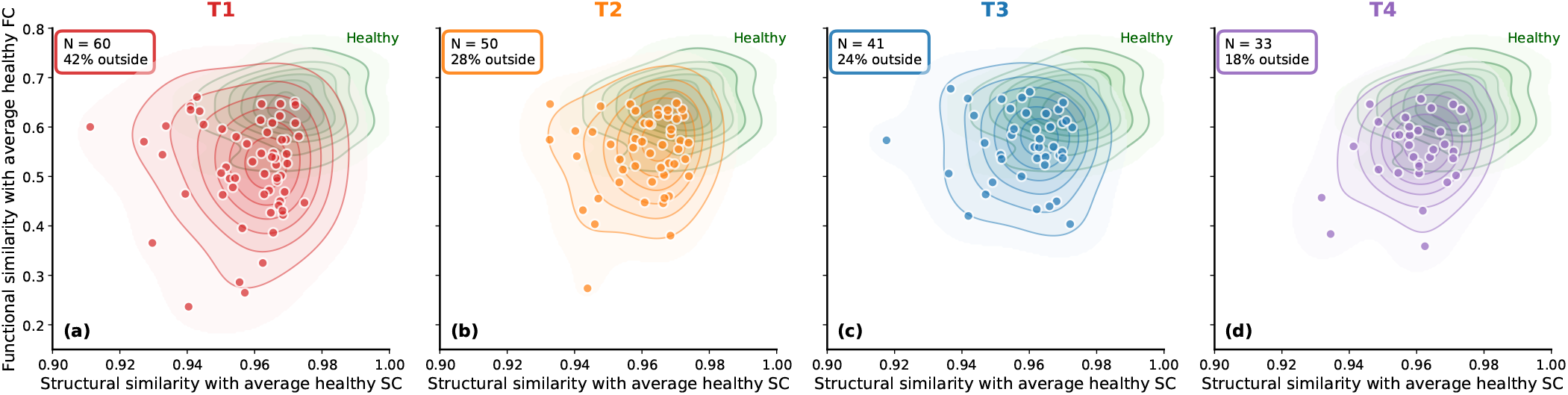
Longitudinal convergence of the post-stroke connectome toward the healthy manifold. (a–d) Each panel shows a bivariate embedding of stroke patients at acute (*T*_1_), early sub-acute (*T*_2_), late sub-acute (*T*_3_), and chronic (*T*_4_) stages. Each point denotes one patient positioned by their similarity (Pearson correlation) to a healthy reference template derived from the ECONS cohort, computed separately for structural connectivity (SC; x-axis) and functional connectivity (FC; y-axis). The shaded density indicates the distribution of healthy controls and patients and their 95% confidence, defining the range of (healthy) normative variability. Percentages report the fraction of patients outside this healthy envelope at each stage. Patients show a pronounced structure–function dissociation early after stroke, with relatively preserved SC similarity but reduced and highly variable FC similarity. Across the first year, patient trajectories shift toward the healthy density peak primarily along the FC axis, whereas SC similarity remains comparatively stable, consistent with recovery dominated by functional reconfiguration on top of a largely invariant structural scaffold. Substantial inter-individual dispersion persists at late stages, reflecting heterogeneous network-level consequences of injury and compensatory adaptations.

In this space, the acute stroke acted as a strong perturbation that scattered individuals away from the healthy manifold. At *T*_1_, 42% of patients lay outside the healthy template, driven primarily by reduced FC similarity despite comparatively preserved whole-brain SC similarity. This structure–function dissociation is consistent with connectional diaschisis, in which focal lesions induce widespread remote functional consequences through disconnection and network-level rebalancing [44]. More generally, it is in line with prior work showing that anatomy constrains, but does not uniquely determine, functional coupling, which can be reshaped through indirect pathways and distributed interactions across the connectome [84, 85].

During their first year post-stroke, patients showed a systematic and individualized drift back to the healthy manifold, reflected by a progressive reduction in outliers. Importantly, this shift was not driven by global “structural repair”. Structural similarity remained comparatively stable over time, consistent with the largely stationary structural backbone reported in Fig. 3**a**, whereas functional similarity increased monotonically. This pattern matches models in which the structural connectome provides relatively rigid constraints, while functional interactions remain more flexible and can reorganize over months as network dynamics settle into new stable configurations [81, 82].

Convergence toward the healthy manifold was nonetheless incomplete. Substantial dispersion persisted at every stage, consistent with heterogeneity in lesion anatomy, compensatory strategies, and the distributed network consequences of injury [50, 86, 87]. This state-space view provides a compact whole-brain account of recovery. Rather than simply reverting to a generic template, the post-stroke brain appears to navigate a constrained dynamical landscape that increasingly overlaps with the healthy structure–function manifold, while retaining marked individual variability shaped by the injury.

### Behavioral recovery is domain-specific and only partly explained by lesion size

Having established that post-stroke recovery involves both stable global identity and ongoing network remodeling, we next asked how these connectome changes relate to clinical recovery outcomes. We first summarized the longitudinal behavioral profile of the cohort using low-dimensional-composite scores derived for each behavioral domain from the full neuropsychological battery via non-negative matrix factorization (NMF), yielding single domain scores comparable across time points (Fig. 6). In this representation, higher values indicate *greater* impairment. Across the first year, most domains showed a clear reduction in impairment from *T*_1_ to *T*_4_, consistent with overall behavioral improvement. Importantly, recovery was not uniform across domains. Motor and sensory scores were already low at *T*_1_ and remained low across time, with comparatively little dispersion across individuals. In contrast, higher-order cognitive domains such as attention and executive function showed substantial inter-individual variability, indicating a wide range of recovery phenotypes.

**FIG. 6.**
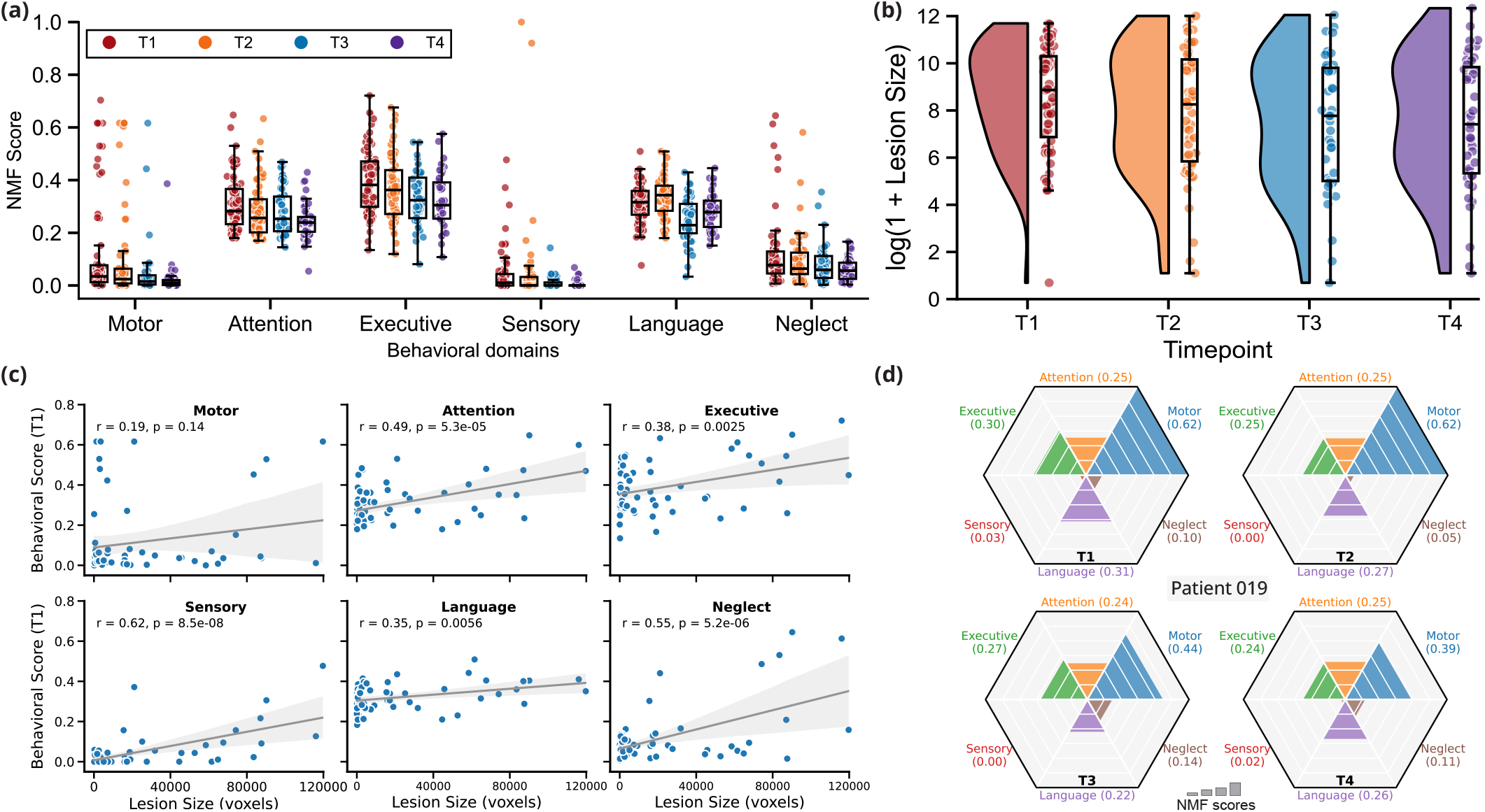
Domain-specific behavioral recovery and its relationship to structural lesion burden. **(a)** Behavioral composite scores across six domains (Motor, Attention, Executive, Sensory, Language, Neglect) over time. Scores were derived using Non-negative Matrix Factorization (NMF) to ensure cross-session comparability; higher scores indicate greater clinical impairment. Boxplots show median, interquartile range, and 1.5 × IQR whiskers at each timepoint (T1: ∼1 week, T2: ∼ 3 weeks, T3: ∼ 3 months, T4: ∼ 1 year) **(b)** Distribution of structural lesion burden across the cohort, quantified as the log-transformed number of voxels (log(1 + voxels)). **(c)** Correlation between acute lesion volume and behavioral impairment at T1. While most domains (Attention, Executive, Sensory, Language, Neglect) show a significant positive correlation with total lesion size, motor impairment exhibits a notable dissociation (*r* = 0.19, *p* = 0.14), suggesting that motor outcomes are driven more by lesion topography than gross volume. **(d)** Example of longitudinal profile (patient P019) showing domain scores across time points, illustrating within-subject heterogeneity and delayed improvement in motor impairment emerging from T3 onward. Parentheses report the NMF domain composite score (arbitrary units; scaled to [0,1]) at each timepoint; higher values indicate greater impairment. This individual example is shown for illustration and is not intended to be representative of the full cohort.

We next examined the extent to which early impairment could be attributed to the sheer scale of the structural damage. Lesion burden, quantified as log-transformed lesion volume (Fig. 6**b**), was correlated with acute composite scores at *T*_1_ (Fig. 6**c**). While most behavioral domains showed significant associations with lesion size, indicating that larger lesions typically produce more severe acute deficits, motor impairment was a notable exception. The weak dependence of motor scores on lesion volume aligns with the established principle that motor outcome is governed less by overall lesion size than by whether the lesion disrupts critical descending motor pathways, such as the corticospinal tract [88, 89].

To further illustrate the granularity of individual trajectories, we highlight the longitudinal profile of one representative case (patient P019; Fig. 6**d**). This patient showed severe acute motor impairment, followed by a delayed improvement that became apparent from *T*_3_ onward, underscoring that recovery can unfold on distinct timescales within the same individual and across domains.

### Multivariate brain–behavior coupling reveals a predictive acute functional signature

The behavioral trajectories across the first post-stroke year revealed both a shared recovery trend and substantial inter-individual variability, motivating a direct test of whether early connectome organization carries prognostic information about later clinical status. We therefore implemented a two-stage brain–behavior framework that combines multivariate association with out-of-sample prediction (Fig. 7). In the first stage, we used Partial Least Squares Correlation (PLSC) to identify latent modes that maximize covariance between whole-brain functional connectivity (brain block) and the set of behavioral domain scores (behavior block) [75]. In the second stage, we converted the significant PLSC brain component into an interpretable connectivity mask and used this reduced feature set to constrain ridge regression models, evaluated with leave-one-subject-out (LOSO) cross-validation. In a way, this strategy follows connectome-based prediction principles [27], while explicitly leveraging multivariate brain–behavior structure. To avoid data leakage [76], we enforced strict temporal separation between feature discovery and prediction. Behavioral scores were first residualized to remove linear time effects, ensuring that predictions were not driven by simple mean recovery trends. Furthermore, PLSC feature selection was performed solely on *T*_*i*_ data, and the resulting connectivity mask was then applied to predict behavior at strictly independent follow-up time points (i.e. *T*_*i*+1_ to *T*_4_).

**FIG. 7.**
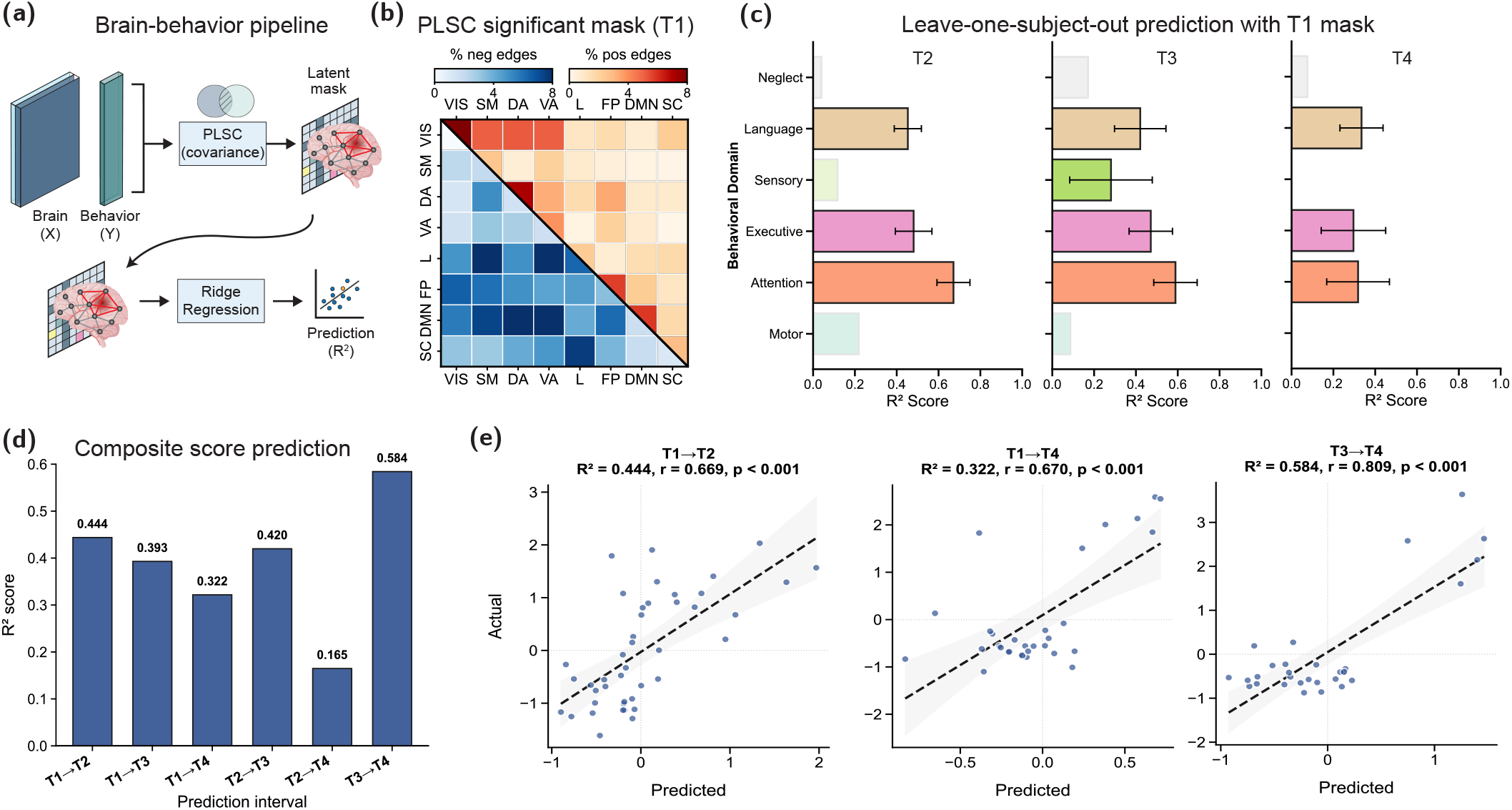
Multivariate brain–behavior mapping and longitudinal forecasting of clinical outcomes. **(a)** Two-stage predictive framework coupling Partial Least Squares Correlation (PLSC) with Ridge regression. Latent axes of maximal covariance between functional connectivity (Brain *X*) and behavioral scores (Behavior *Y* ) are identified via PLSC. Significant components generate a connectivity mask for Ridge regression, validated using a leave-one-subject-out (LOSO) approach to predict outcomes at subsequent time points (*T*_*i*_ → *T*_*i*+1…4_). **(b)** Network-level aggregation of the significant PLSC connectivity mask derived at the acute stage (*T*_1_). Heatmaps display the percentage of significantly negative (blue) and positive (red) edges, highlighting the prominent contribution of higher-order association systems (FP, DMN) and subcortical structures (SC) to the acute behavioral phenotype. **(c)** LOSO prediction performance (*R*^2^) for individual behavioral domains using the *T*_1_ latent mask. The model successfully forecasts language, executive, and attention outcomes at follow-up (*T*_2_, *T*_3_, *T*_4_), whereas motor and neglect domains show negligible predictability from this functional signature. **(d)** Prediction accuracy for a global composite behavioral score across longitudinal intervals. Accuracy decays as the temporal gap from *T*_1_ increases but peaks during the late sub-acute to chronic transition (*T*_3_ → *T*_4_; *R*^2^ = 0.584), reflecting late-stage stabilization of the brain–behavior relationship. **(e)** Scatter plots correlating actual versus predicted global scores for some of the intervals reported in panel (d).

At the acute stage, PLSC identified a robust multi-variate connectivity pattern with significant edges distributed across canonical systems and carrying both positive and negative weights (Fig. 7**b**). When summarized at the network level, the latent mask emphasized between-network interactions and a characteristic balance of positively and negatively weighted edges, providing an interpretable functional signature of acute impairment that could be propagated forward as a fixed feature set. Using this acute PLSC-derived mask, ridge regression yielded selective, domain-specific prediction at follow-up (Fig. 7**c**). In LOSO prediction, acute connectome features explained meaningful variance in language, executive, and attention outcomes across subsequent stages, whereas other domains showed limited or no predictability from the same mask. This specificity was not trivially explained by model flexibility. Prediction performance was stable across ridge regularization strengths (main analyses: *α* = 1), and models trained on the full, unmasked FC did not yield significant predictions across hyperparameter choices. These results indicate that the covariance-informed mask isolates a subset of altered connections that is disproportionately informative for later cognitive outcomes.

We next assessed whether the same framework captures recovery at a global level. To do so, we derived a single composite impairment score per patient and time point by summarizing the joint domain profile (convex-hull composite; see Methods for details), and repeated the PLSC-mask and ridge prediction across longitudinal intervals (Fig. 7**d–e**). Prediction accuracy decreased as the temporal gap from baseline increased, consistent with the accumulating influence of ongoing reorganization and intercurrent clinical factors over longer horizons. Notably, prediction was strongest for the late interval *T*_3_ → *T*_4_ (*R*^2^ = 0.584, *r* = 0.809, *p <* 0.001), exceeding performance for *T*_1_ → *T*_2_ (*R*^2^ = 0.444, *r* = 0.669, *p <* 0.001) and *T*_1_ → *T*_4_ (*R*^2^ = 0.322, *r* = 0.670, *p <* 0.001). This late-stage peak suggests that once patients reach the late sub-acute phase, behavior and the underlying functional configuration become more tightly coupled and therefore more predictable.

Together, these analyses link early connectome organization to later clinical status in a longitudinally separated pipeline. PLSC isolates a multivariate acute connectivity signature [75], and ridge regression shows that this signature predicts recovery outcomes in specific cognitive domains as well as in a global composite measure.

## DISCUSSION

Stroke is increasingly viewed as a network disorder, in which focal lesions propagate their impact across large-scale systems by inducing disconnection and remote physiological alterations [3, 4, 44, 45, 47, 56]. Leveraging multimodal longitudinal data from the TiMeS cohort [71], we provided a multi-scale view of post-stroke connectome dynamics across the first year of recovery. Despite substantial focal damage, individual functional identity is remarkably resilient, consolidating within the first three weeks into a stable, patient-specific configuration. This early stabilization of global identity does not imply functional stasis, as link-wise connectivity continues to reorganize over months with a distinct temporal ordering of hyper- and hypo-connectivity. At the macroscale, patients show a gradual shift toward the range of healthy structure–function variability, despite a largely stationary structural backbone. Together, these results frame recovery as constrained navigation of a dynamical structure–function landscape, in which a stable individual scaffold emerges quickly while selected circuits continue to remodel within anatomical limits. Crucially, this early network organization is directly linked to clinical recovery, forming a stable foundation that predicts long-term behavioral outcomes.

### The paradox of rapid stabilization and persistent plasticity

Connectome fingerprinting was originally established as the capacity of functional connectivity to identify healthy individuals across sessions and conditions [15, 17, 28, 31, 90, 91]. Our data extend this framework to stroke, demonstrating that a focal injury does not permanently destabilize the idiosyncratic architecture of the individual brain. Instead, the connectome consolidates rapidly: by the early sub-acute phase (3 weeks), patients are as identifiable to themselves as they are in the late subacute/chronic phase. This suggests that the post-stroke brain does not remain in prolonged flux; rather, it quickly relaxes into a new “recovery configuration”, consistent with dynamical systems perspectives where brain activity occupies a restricted repertoire of metastable states constrained by the underlying anatomy [81, 82]. In this view, stroke acts as a high-energy perturbation that displaces the system from its pre-injury basin of attraction, but the system rapidly settles into a new local minimum rather than drifting without structure.

Crucially, this global stability coexists with—and perhaps relies on—the plasticity of specific functional anchors. We found that higher-order association networks (DMN, FPN) maintained the strongest longitudinal stability, whereas subcortical and sensorimotor loops were more variable. This aligns with the notion that association cortices provide an integrative backbone for cognition [36, 78], while cortico-subcortical loops—which are particularly vulnerable to lesion-induced disruption—undergo more pronounced reweighting as sensorimotor control is re-optimized. This network-specific stability helps reconcile why global identifiability can plateau early even as clinically meaningful recovery continues: a stable functional identity provides the necessary background for the targeted remodeling of specific circuits.

### Decoupling of structural injury and functional reorganization

A central contribution of this work is to separate stable from dynamic components of post-stroke network change. The structural disconnection backbone estimated from diffusion data was largely stable across time, consistent with the view that the anatomical consequences of infarction stabilize early and show only limited changes further on, even when their network consequences are widespread [45, 47]. Functional connectivity, in contrast, continued to reorganize well into the chronic phase, consistent with longitudinal work showing evolving disruptions and partial normalization over months [35, 37, 38, 49]. This dissociation echoes a broader principle that anatomy constrains but does not uniquely determine function, which is shaped by indirect pathways, distributed interactions, and state-dependent dynamics [81, 82, 84]. It also provides a network-level substrate for diaschisis, where focal damage produces remote dysfunction through disconnection and rebalancing of interactions across systems [44].

Beyond this structure–function decoupling, our results refine the temporal interpretation of post-stroke plasticity. Specifically, we initially observed an early, transient increase in functional hyper-connectivity. This acute response was followed by a delayed rise in hypoconnectivity that peaked months post-stroke, disproportionately affecting association networks and cortico-subcortical interactions. Early hyper-connectivity may reflect compensatory recruitment of available resources and short-term stabilization of communication, consistent with reports of distributed engagement in early recovery [40, 62, 65]. However, persistent hyperconnectivity is not necessarily adaptive. Models of interhemispheric imbalance and maladaptive contralesional influence predict poorer outcomes when compensatory recruitment becomes entrenched [64], especially in well-recovered patients with limited remaining deficits [92]. The later emergence of widespread hypo-connectivity, especially in higher-order association systems and subcortical interactions, is consistent with network-disconnection accounts of cognitive deficits after stroke [45, 46, 56]. Together, these findings argue against a simple narrative of monotonic “normalization”. Instead, recovery appears to unfold through sequential phases of coupling and decoupling as the system rebalances integration and segregation and settles into a new stable configuration to limit remaining deficits [49].

### Recovery as a trajectory in a structure–function manifold

Patients were displaced acutely away from the healthy structure–function regime, mainly along the functional axis, and then gradually shifted back toward the envelope of healthy variability. This picture aligns with whole-brain modeling views in which the structural connectome shapes an attractor-like landscape of feasible functional states [82, 83]. In this framing, the stationary structural coordinate reflects the rigidity of the anatomical scaffold after focal injury, whereas the gradual increase in functional similarity reflects re-tuning of dynamics constrained by surviving pathways. Convergence was nonetheless incomplete. Substantial dispersion persisted across stage, consistent with variability in lesion topography, disconnection profiles, and compensatory strategies [45, 47, 87]. Rather than noise, this dispersion may reflect distinct recovery phenotypes in which injury reshapes the structure–function mapping differently, leading some patients to stable but suboptimal configurations.

### Multivariate brain–behavior signatures of recovery

A practical motivation for individualized network mapping is prognosis, as it is still a challenge based on clinical evaluation only. Using domain-specific behavioral composites that are comparable across time [43], we found that while acute impairment in most domains scaled with overall lesion volume, motor deficits exhibited a notable dissociation. This matches a well-established principle that motor impairment and recovery depend strongly on lesion topography and corticospinal tract integrity rather than lesion size alone [43, 58, 88, 89, 93]. However, it is important to note that the limited variability of motor scores within our specific cohort may have statistically attenuated this association. The interplay of these biological constraints and potential analytic artifacts aligns closely with ongoing debates surrounding proportional recovery rules [57, 94–96].

Our brain–behavior analyses extend lesion-based accounts by directly coupling distributed functional architecture to multidomain behavior. PLSC is well-suited for this setting because it extracts latent modes that maximize covariance between high-dimensional brain features and multivariate behavior [75]. Using PLSC-derived masks to constrain ridge regression, we found that acute functional signatures carried prognostic information for language, executive function, and attention. Notably, these are domains that depend on distributed association networks and long-range integration [46, 56]. In contrast, motor and neglect outcomes were less predictable from the same functional mask. While this aligns with the tighter dependence of motor outcomes on specific descending pathways and the clinical heterogeneity of spatial-attention deficits, this limited predictability must be interpreted with caution. As noted above, the restricted variability of motor scores within our specific cohort likely attenuated the performance of these statistical models. Methodologically, our temporally separated feature discovery and follow-up prediction design follows best practices in connectome-based predictive modeling and addresses known pitfalls of leakage and overly optimistic estimates in moderate samples [27, 76]. The peak predictability from late sub-acute to chronic stages further suggests that after ∼ 3 months, the coupling between functional architecture and clinical phenotype becomes tighter and more stable, consistent with consolidation of a chronic recovery configuration.

### Limitations and future directions

Several limitations should guide interpretation. Missing modalities across time points reduce balanced sampling, and replication in larger cohorts with denser follow-up will be important. Resting-state fMRI after stroke can be influenced by vigilance, medication, and vascular or hemodynamic alterations, and future work should incorporate physiological monitoring and measures of vascular reactivity to better dissociate neuronal from vascular contributions. Predictive modeling in moderate samples remains vulnerable to instability, and external validation and multi-site generalization tests are critical, as emphasized by the broader neuroimaging prediction literature [76]. Extending fingerprinting and prediction to multimodal signals that show robust identifiability and may be more scalable clinically could strengthen translation [20, 21, 23].

Finally, functional connectivity and downstream inference can be sensitive to demographic and clinical covariates, as well as to how confounds are handled in multivariate pipelines [77, 97–99]. Although the main text reports results without explicitly residualizing age, sex, or lesion burden from FC, we consider robustness by re-estimating key effects after residualizing FC using linear mixed-effects models that included these covariates (Methods). This alternative specification yielded quite similar and concordant results for fingerprint stabilization, network-level hyper-/hypo-connectivity trajectories, and brain–behavior associations (Supplementary Figs. S6, S7, S8), supporting the interpretation that our main conclusions reflect genuine longitudinal reconfiguration rather than bein driven by these confounders.

In conclusion, our findings highlight a critical dissociation in post-stroke recovery: while structural damage remains largely stable, the brain rapidly consolidates a new, stable functional identity. This early stabilization provides a crucial macro-scale architecture, which allows specific networks to reorganize over the following months. Ultimately, mapping these dynamic functional shifts offers a powerful and individualized framework for predicting long-term behavioral recovery.

## METHODS

### Participants and study design

We analyzed longitudinal data from the TiMeS (Toward individualized medicine in stroke) cohort [71], a study examining stroke recovery with multimodal neuroimaging and detailed neuropsychological evaluation. Patients were assessed at four post-stroke stages: acute (*T*_1_, ∼ 1 week), early sub-acute (*T*_2_, ∼ 3 weeks), late subacute (*T*_3_, ∼ 3 months), and chronic (*T*_4_, ∼ 12 months). The present analyses included up to *N* = 64 stroke patients (availability varied across time points and modalities; mean age 66.5 ± 13.9 years; 18 females). See Table I for exact sample sizes per time point and analysis. Inclusion criteria were *(1)* age ≥ 18 years; *(2)* diagnosis of a first-ever or recurrent, ischemic or hemorrhagic stroke; *(3)* enrollment within 7 days of the stroke incident; *(4)* presence of upper-limb motor impairment, objectified by clinical assessment. For the full eligibility criteria, refer to [71]. Two healthy control cohorts were used to define normative reference distributions, ECONS (*N* = 31, mean age 69.1 ± 4.2) and TrainStim (*N* = 10, mean age 69.9 ± 4.6) [100, 101], acquired with harmonized protocols and processed with the same pipeline as the patient data. ECONS served as the primary reference for computing clinical identifiability and functional deviation statistics. All participants provided written informed consent in accordance with the Declaration of Helsinki, and the study was approved by the local Ethics Committee (Canton of Valais and Canton of Geneva, Switzerland). Because longitudinal follow-up was incomplete for some participants, each analysis used all available data for the required modality and time point unless stated otherwise. Analyses requiring joint structure–function information were restricted to participants with complete multimodal data for the corresponding sessions.

**TABLE I.** Data availability across the TiMeS longitudinal cohort. Entries report the number of stroke patients with available data for resting-state fMRI (BOLD), BOLD plus behavioral assessment, BOLD plus lesion information, or all modalities jointly, at each post-stroke session. Columns labeled *T*_*i&*_*T*_*j*_ indicate the number of patients with data available at both time points, enabling paired longitudinal analyses. Columns labeled T1–T3 and T1–T4 indicate the number of patients with complete longitudinal coverage across the corresponding interval.

| | T1 | T2 | T3 | T4 | $T1\&T2$ | $T1\&T3$ | $T1\&T4$ | $T2\&T3$ | $T2\&T4$ | $T3\&T4$ | T1-T3 | T1-T4 |
| --- | --- | --- | --- | --- | --- | --- | --- | --- | --- | --- | --- | --- |
| **BOLD** | 62 | 53 | 44 | 40 | 45 | 39 | 33 | 37 | 31 | 31 | 32 | 22 |
| **BOLD & Behavioral** | 56 | 53 | 43 | 39 | 43 | 37 | 31 | 36 | 30 | 30 | 31 | 21 |
| **BOLD & Lesions** | 60 | 50 | 41 | 33 | 43 | 37 | 27 | 33 | 24 | 25 | 30 | 17 |
| **All** | 54 | 50 | 40 | 33 | 41 | 35 | 26 | 32 | 24 | 25 | 29 | 17 |

### MRI acquisition and preprocessing

Magnetic resonance imaging (MRI) data were acquired on a 3T Siemens Prisma scanner. Resting-state BOLD fMRI was acquired using a multi-band echo-planar imaging (EPI) sequence (TR = 1,250 ms; TE = 32 ms; flip angle = 58^°^; 2 mm isotropic resolution; multiband factor 5). Each session comprised 8.13 minutes of eyes-open (375 volumes). Structural and diffusion-weighted imaging (DWI) sequences were acquired at each session using the parameters listed in the TiMeS protocol [71]. Resting-state BOLD data were preprocessed using a standard pipeline including slice-timing correction, realignment, and co-registration to the structural scan. To mitigate non-neural noise, we applied nuisance regression (6 motion parameters, mean white matter, and cerebrospinal fluid signals) and bandpass filtering (0.01–0.15 Hz).

#### Diffusion preprocessing and structural connectome estimation

Diffusion-weighted data underwent the following pre-processing steps using MRtrix3 (version 3.0.4; https://www.mrtrix.org/) and FSL (version 6.0.6.4; https://fsl.fmrib.ox.ac.uk/fsl/docs/): Gibbs ringing was removed with MRtrix3, brain extraction was performed using FSL BET, susceptibility-induced distortion correction was applied with FSL topup, eddy-current and motion corrections were implemented using FSL eddy, and bias-field inhomogeneity was corrected with FSL FAST. Segmentation, including the brainstem, was performed using FreeSurfer (version 7.4; https://surfer.nmr.mgh.harvard.edu/). Anatomical images were registered to diffusion space using ANTs (version 2.4.4; https://github.com/ANTsX/ANTs) to ensure alignment, and tissue-type maps were generated for multi-tissue modeling. Fiber orientation distributions were estimated using multi-shell, multi-tissue constrained spherical deconvolution (MSMT-CSD) [102], and whole-brain tractography was performed using probabilistic tracking with 10 million streamlines. Streamline weights were refined using SIFT2 [103] to improve the biological accuracy and reduce the weighting of false positive connections. A subject-specific structural connectome was obtained by mapping streamline weights to the Glasser atlas regions using the MRtrix3 tck2connectome function.

#### Functional and structural connectome construction

For each session, whole-brain functional connectivity (FC) was estimated using a combined cortical–subcortical parcellation of 377 regions: 360 cortical parcels from the Glasser parcellation atlas [72] and 17 subcortical as provided by the HCP release (filename “Atlas ROI2.nii.gz”; left and right: thalamus, caudate, putamen, pallidum, amygdala, accumbens, ventral diencephalon; and brainstem). Hippocampal parcels of the Glasser parcellation were considered as subcortical ones. Individual functional connectomes were constructed by computing the Pearson correlation coefficient between all pairs of regional time series, resulting in a 377 × 377 symmetric association matrix. For network-level analyses, nodes were assigned to seven canonical Resting State Networks (RSNs) [78] (Visual, Somatomotor, Dorsal Attention, Ventral Attention, Limbic, Frontoparietal, Default Mode) plus Subcortical structures.

#### Connectome fingerprinting and identifiability metrics

To quantify the stability of patient-specific functional architecture, we employed the connectome fingerprinting framework [15, 17]. For subject *s* at time points *T*_*i*_ and *T*_*j*_, longitudinal self-similarity was defined as

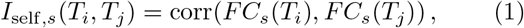

For each interval (*T*_*i*_, *T*_*j*_), we constructed an identifiability matrix by correlating vectorized FC patterns across all pairs of subjects (test at *T*_*i*_, retest at *T*_*j*_). Within-subject similarity corresponds to diagonal entries, whereas between-subject similarity corresponds to off-diagonal entries. Differential identifiability was defined as

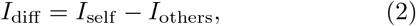

capturing whether, on average, subjects remain more similar to themselves than to other individuals.

#### Clinical similarity to healthy controls

To quantify deviation from healthy norms, we computed an individual clinical similarity index adapted from Ref. [28]. For each stroke subject *s* at time *T*_*i*_,

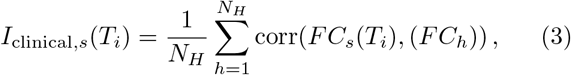

where *FC*_*h*_ denotes the FC of healthy individual *h* and *N*_*H*_ is the number of controls.

#### Statistical inference for longitudinal fingerprinting

To account for repeated measures and unbalanced sampling, we tested changes in *I*_self_ across longitudinal intervals using linear mixed-effects models (LMMs) with subject-specific random intercepts:

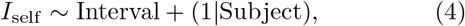

where Interval is a categorical fixed effect representing the time pairing (e.g., *T*_1_–*T*_4_, *T*_2_–*T*_4_, *T*_3_–*T*_4_).Models were fit using restricted maximum likelihood (REML), and inference on fixed effects was performed using Wald tests (FDR-corrected). Effect sizes were summarized using Cohen’s *d* [80]. Mann-Whitney U tests were used for *I*_*clinical*_ comparisons, with FDR correction at *q <* 0.05.

#### Single-edge fingerprinting and link reliability

To localize stable connections that support individual identifiability, we estimated the intraclass correlation coefficient (ICC) for each edge using a one-way random-effects model [74]. For a given interval (*T*_*i*_, *T*_*j*_), each subject contributed two observations per edge, *e*_*n*_(*T*_*i*_) and *e*_*n*_(*T*_*j*_). ICC was computed as

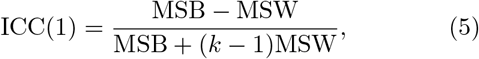

where MSB and MSW denote between-subject and within-subject mean squares, respectively, and *k* = 2 observations per subject. Edges with ICC *>* 0.6 were defined as reliable *individual edges*, consistent with thresholds used in prior fingerprinting work [23, 31]. To mitigate sample-size dependence of ICC estimates, we used bootstrapping: for each interval, ICC matrices were computed over 200 bootstrap resamples using a fixed number of subjects (*N* = 15) drawn from those available for that interval, and then averaged. Node-level contributions were obtained by summing ICC values across incident edges, yielding a nodal ICC score that reflects the extent to which a node’s connectivity profile contributes to identifiability.

#### Structural disconnection and disconnectome estimation

To characterize the structural backbone of stroke, we estimated a disconnectome from diffusion-derived structural connectivity by overlaying each participant’s binary lesion mask onto the tractogram and retaining only streamlines intersecting the lesion. For each subject and time point, we derived an indicator of structurally disrupted edges by thresholding edges with non-zero disconnection evidence in the disconnectome matrices. Group summaries were computed as the fraction of subjects exhibiting a disrupted edge within each within- and between-network block. Robustness was assessed by comparing the primary disconnectome estimation against alternative lesion-based procedures, including BCB toolkit-based disconnection [79], yielding convergent group-level patterns (Supplementary Figs. S5).

#### Whole-brain structure–function state-space embedding

To provide a compact description of longitudinal brainstate shifts, we embedded each patient session into a joint structure–function space defined by similarity to a normative healthy template. Structural similarity was computed as cosine similarity between the vectorized subject structural connectome and the mean healthy structural connectome (ECONS). Functional similarity was computed analogously using the vectorized FC and the mean healthy FC. The healthy reference distribution was represented with a two-dimensional Gaussian kernel density estimate (KDE), and a normative envelope was defined by thresholding this density at 95%. Patient sessions falling outside this envelope were classified as outside healthy variability. Longitudinal changes were summarized by a drift toward the peak of the healthy density and by the time-dependent fraction of outliers.

#### Behavioral scores

At each time point, patients underwent a multidomain behavioral assessment comprising 40 validated tests spanning motor, sensory, and cognitive functions [71]. Missing data were imputed using principal component–based methods (missMDA). Motor and sensory scores were normalized as ratios between affected and unaffected limbs, and all measures were subsequently min–max scaled such that higher values reflected greater impairment. To derive a single interpretable score per domain, variables with ceiling effects or high intercorrelations (*r >* 0.8) within the same domain were excluded. Non-negative matrix factorization (NMF) [104] was then applied at T1 to extract one latent, non-negative feature per domain, yielding normalized (0–1), additive measures of impairment across Motor, Attention, Executive, Language, and Neglect domains, with higher values consistently reflecting greater impairment. The resulting model was applied to subsequent timepoints to ensure consistent scaling and enable longitudinal comparisons. Model performance was assessed using variance accounted for (VAF), with the derived features explaining 66% of the variance for Neglect and greater than 80% for all other domains. Lesion size was quantified as the number of lesioned voxels at *T* 1 and analyzed as log(1 + voxels). Associations between lesion volume and acute behavioral impairment were assessed using Spearman correlation.

#### Detrending behavioral scores for longitudinal prediction

Because behavioral scores exhibited systematic time dependence, we reduced temporal confounding before prediction. For subjects with complete longitudinal behavioral data, we fit an ordinary least squares model for each domain as a function of time (coded as 0, 1, 2, 3 for *T*_1_ … *T*_4_) and retained residuals as detrended behavioral scores. For subjects without complete longitudinal behavioral data, we retained raw scores to avoid extrapolation. This procedure reduces apparent predictability driven by shared temporal trends rather than brain-behavior coupling.

#### Multivariate brain–behavior association and leakage-aware prediction

We related FC features to multidomain behavior using Partial Least Squares Correlation (PLSC) [75]. At each time point *T*_*i*_, we assembled a brain matrix *X* (subjects × edges, using vectorized FC) and a behavioral matrix *Y* (subjects × domains, using detrended NMF scores). PLSC was computed with `pyls` (`behavioral_pls`) Python package [105], assessing latent-variable significance via permutation testing (*n*_perm_ = 1,000) and edge reliability via bootstrapping (*n*_boot_ = 1,000). Reliable edges were defined using a bootstrap-ratio threshold |BSR| ≥ 2.0.

To test prognostic value while minimizing data leakage, we enforced temporal separation between feature discovery and prediction. Specifically, PLSC was fit only at time *T*_*i*_ using brain and behavior from *T*_*i*_, yielding a latent connectivity mask. This fixed mask was then applied to FC at later sessions (*T*_*i*+1_ … *T*_4_) to predict follow-up behavior. Prediction used ridge regression (L2-regularized linear regression) evaluated with leave-one-subject-out (LOSO) cross-validation. The ridge penalty was set to *α* = 1 in primary analyses, with stable results across alternative values (e.g. *α* = 0.1, 10). Performance was quantified using out-of-sample *R*^2^ for each behavioral domain and for a global composite impairment score.

#### Covariate handling and robustness analyses

Main results are presented using minimally adjusted FC estimates, without explicitly regressing demographic or clinical covariates from the connectomes. To assess whether our findings were sensitive to potential confounding by age, sex, or lesion burden, we repeated the principal analyses after residualizing FC using linear mixed-effects models (LME). Across all core results—fingerprint stabilization, network-level hyper-/hypo-connectivity trajectories, the joint structure– function embedding, and brain–behavior associations— the residualized analyses produced results that were quite similar to the primary analyses (Supplementary Figs. S6-S8).

Covariate correction was implemented independently for each FC edge using a two-stage procedure designed to remove only *population-level* nuisance effects while preserving *subject-specific* connectivity structure (i.e., random intercepts). In the first stage, we estimated age and sex effects using all participants (patients and controls) with the following model:

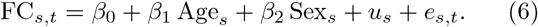

Here, FC_*s,t*_ denotes the FC value for a given edge for subject *s* at session/time point *t*; *β*_0_ is a fixed intercept; *β*_1_ and *β*_2_ are fixed-effect coefficients for age (z-scored) and sex (effect-coded); *u*_*s*_ is the subject-specific random intercept capturing stable between-subject differences; and *e*_*s,t*_ is the residual term capturing within-subject/session variability not explained by the model (statsmodels `MixedLM`, REML). We subtracted only the fixed-effect contribution of age and sex, 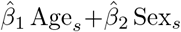 thereby retaining *u*_*s*_ (and thus the subject-specific baseline).

In the second stage, restricted to stroke patients, we modeled the age/sex-adjusted FC values as a function of lesion size:

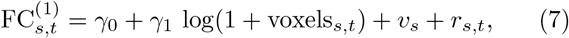

where log(1 + voxels_*s,t*_) is the log-transformed lesion volume (mean-centered within patients), *γ*_0_ and *γ*_1_ are fixed effects, *v*_*s*_ is a patient-specific random intercept, and *r*_*s,t*_ is the residual term. As above, we subtracted only the fixed-effect lesion-size contribution, 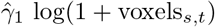, preserving the patient-specific intercept *v*_*s*_.

#### Multiple-comparison correction

Unless stated otherwise, statistical tests were two-sided. Edge-wise analyses controlled false discovery rate across connections using Benjamini–Hochberg correction (*q <* 0.05). Mixed-model contrasts were FDR-corrected across planned comparisons. Effect sizes are reported as Cohen’s *d* [80] where appropriate.

## Code availability

The Python code used in this work will be made available upon acceptance of the manuscript on AS GitHub page.

## Acknowledgment

A.S. has received funding from the European Union’s Horizon 2020 research and innovation programme under the Marie Skłodowska-Curie grant agreement no. 101208090. A.S. thanks D. Orsenigo, M. Nurisso, and M. Diano for helpful discussions. This study was funded by ‘Personalized Health and Related Technologies (PHRT-#2017-205) of the ETH Domain to FCH, the Defitech Foundation (Strike-the-Stroke project, Morges, Switzerland) to FCH, and the Wyss Center for Bio and Neuroengineering (WP030; Geneva, Switzerland) to FCH.

---

**FIG. S1.**
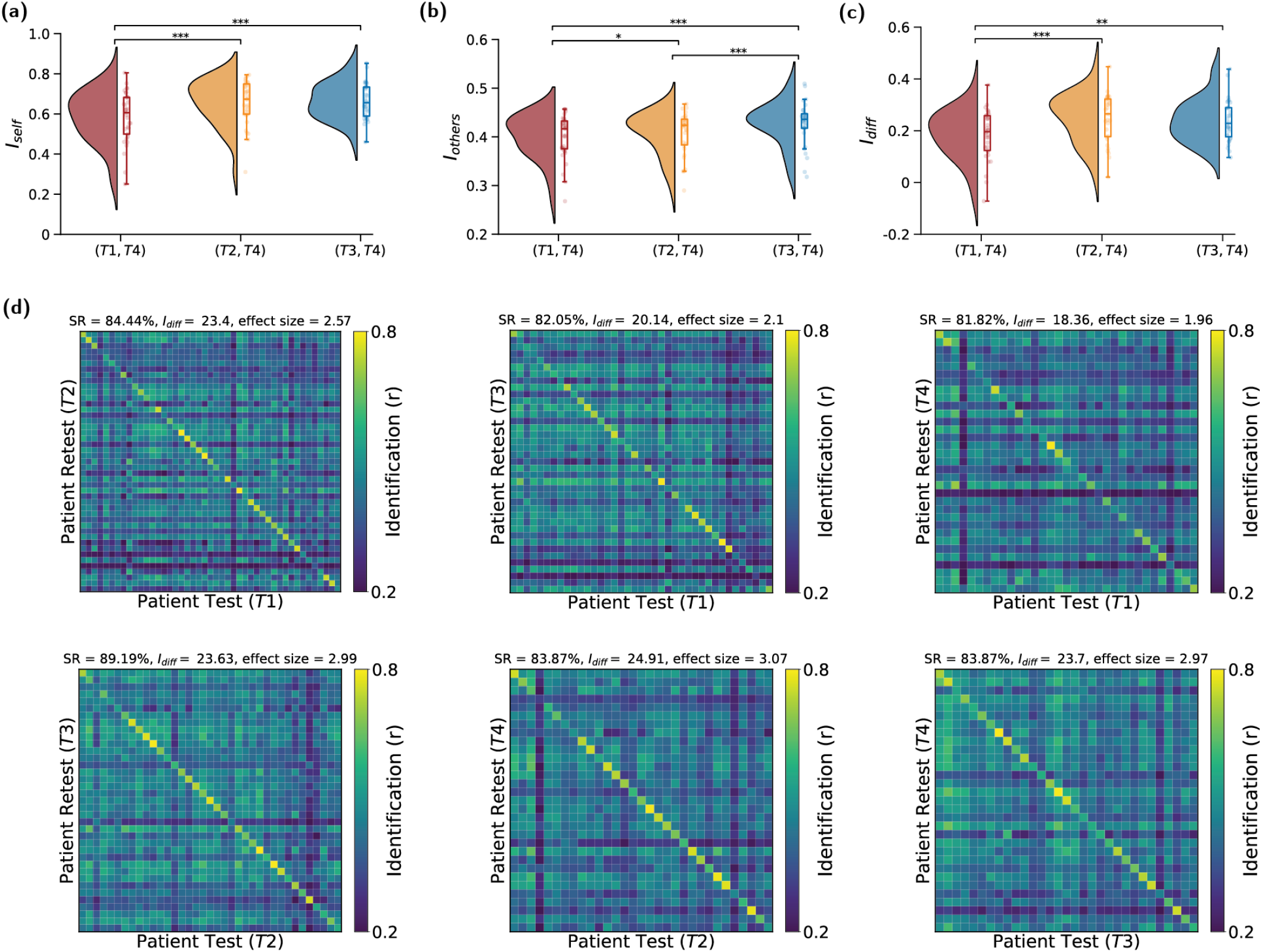
Longitudinal identifiability metrics across all pairs of time points. **(a–c)** Distributions of self-identifiability (*I*_self_ ), between-subject identifiability (*I*_others_), and identifiability difference [*I*_diff_ = (*I*_self_ − *I*_others_]), for connectomes compared against the late-chronic reference (*T*_4_), i.e. between T1–T4, T2–T4 and T3–T4. All three metrics indicate that subject-level reconfiguration is largest between the acute (T1) and early sub-acute (T2) phases, as T1–T4 shows markedly lower identifiability than T2–T4 and T3–T4 (( ∗∗*p <* 0.05, ∗∗*p <* 0.01, Wilcoxon test, FDR-corrected). **(d)** Identifiability matrices for all longitudinal pairings, reporting success rate (SR), mean *I*_diff_, and Cohen’s *d*. Pairings that are closer in time (e.g., *T*_1_–*T*_2_, *T*_2_–*T*_3_) generally show higher SR and larger effect sizes. Notably, despite its shorter temporal separation, *T*_1_–*T*_3_ shows lower SR and smaller effects than later intervals involving *T*_4_ (e.g., *T*_2_–*T*_4_, *T*_3_–*T*_4_), reinforcing the conclusion that a disproportionate reorganization of individual connectome identity is concentrated between the acute (*T*_1_) and early sub-acute (*T*_2_) stages.

**FIG. S2.**
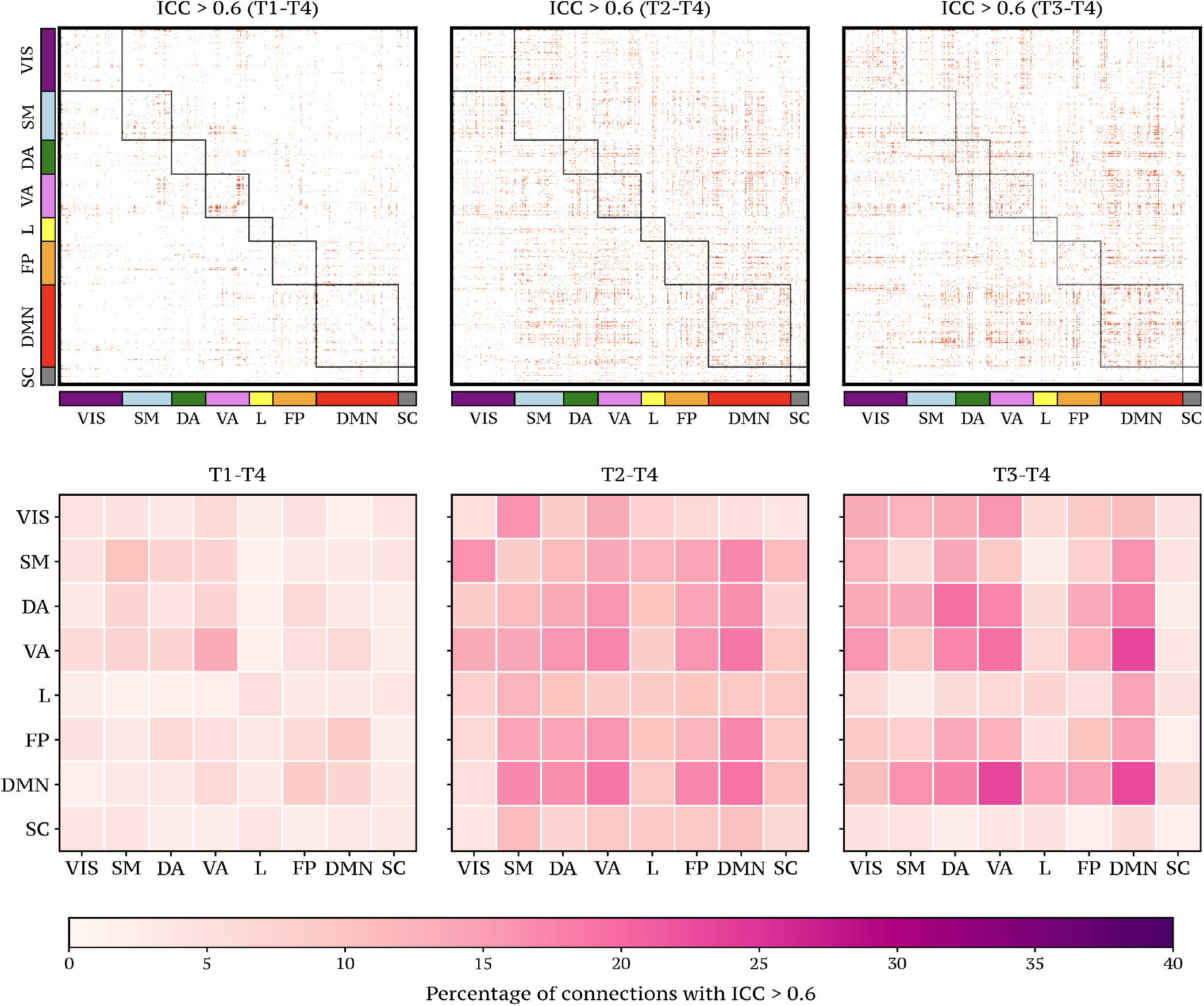
Spatial specificity of connectome fingerprinting across times. Topography of stable individual functional connections, defined by an Intraclass Correlation Coefficient (*ICC*) *>* 0.6, evaluated between early recovery stages (*T*_1_, *T*_2_, *T*_3_) and the chronic baseline (*T*_4_). Matrices display the percentage of these highly stable edges within and between canonical functional networks and subcortical structures. Notably, connections involving higher-order association systems—specifically the frontoparietal (FP) and default mode (DMN) networks—exhibit the highest proportion of stable edges across evaluated intervals, suggesting that these networks form a persistent functional infrastructure that anchors the individual connectome post-stroke. To eliminate variance driven by sample attrition, this analysis was restricted exclusively to the subset of 22 patients common to the analyzed longitudinal intervals. Network abbreviations: VIS, visual; SM, somatomotor; DA, dorsal attention; VA, ventral attention; L, limbic; FP, frontoparietal; DMN, default mode; SC, subcortical.

**FIG. S3.**
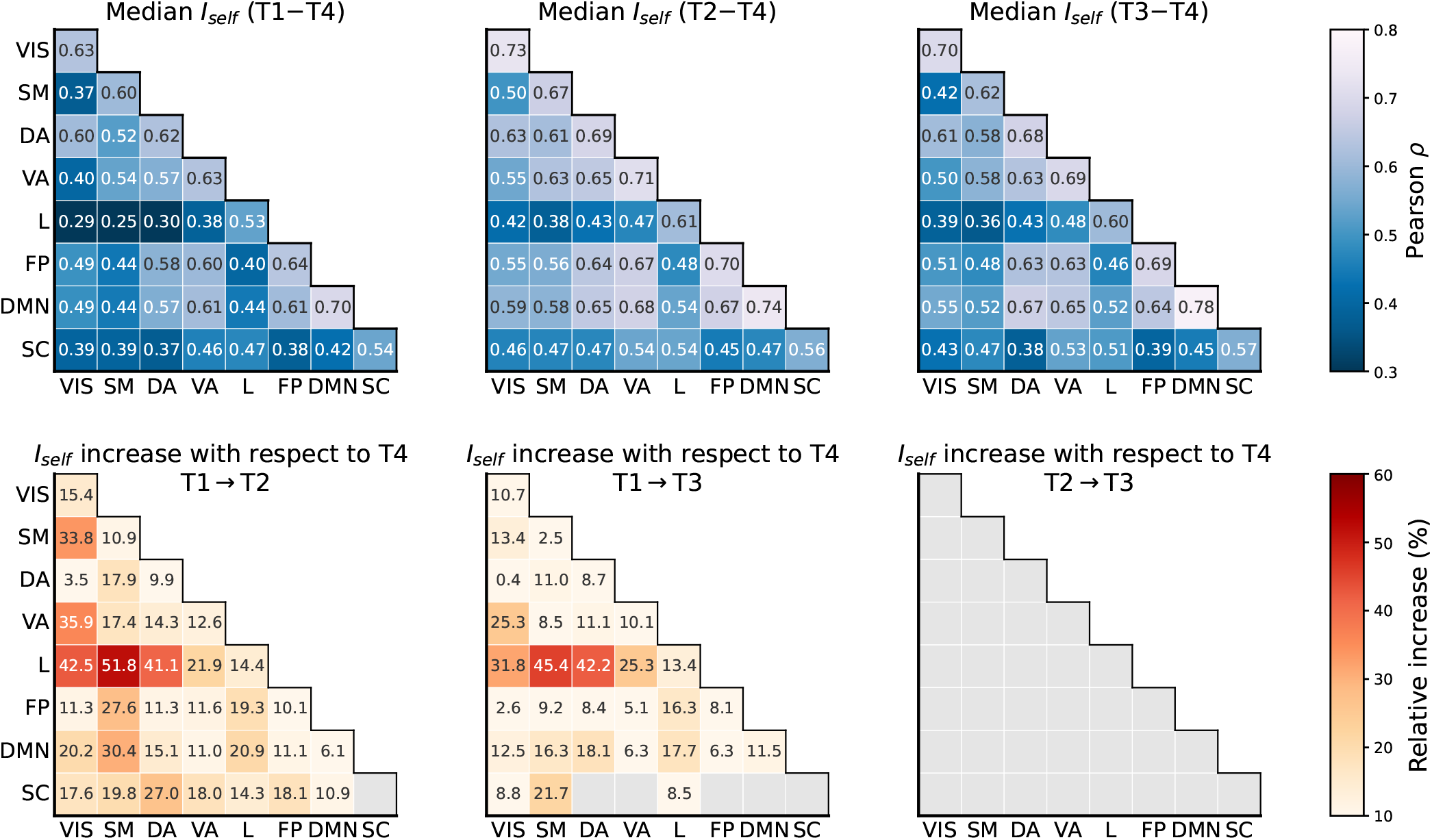
Network-resolved consolidation of longitudinal self-identifiability. Heatmaps report the median within-subject similarity (*I*_self_ ; Pearson correlation) of functional connectivity patterns computed between early sessions and the chronic endpoint (*T*_4_), summarized within and between canonical networks (VIS, SM, DA, VA, L, FP, DMN, SC). Top panels show median *I*_self_ for the (*T*_1_, *T*_4_), (*T*_2_, *T*_4_), and (*T*_3_, *T*_4_) intervals. In line with an early reconfiguration of the individual fingerprint, *I*_self_ increases markedly from *T*_1_ to *T*_2_ (and remains elevated at *T*_3_), whereas differences between *T*_2_ and *T*_3_ are minimal, indicating little additional change after the early sub-acute stage. Bottom panels quantify the relative median increase in *I*_self_ between successive early intervals (e.g., *T*_1_ →*T*_2_, *T*_1_ →*T*_3_, *T*_2_ → *T*_3_), expressed as a percentage of the corresponding *T*_4_-referenced baseline, further emphasizing that the dominant shift occurs between *T*_1_ and *T*_2_ rather than between *T*_2_ and *T*_3_. Statistical significance of interval effects was assessed using linear mixed-effects models with subject-specific random intercepts and FDR correction across network blocks.

**FIG. S4.**
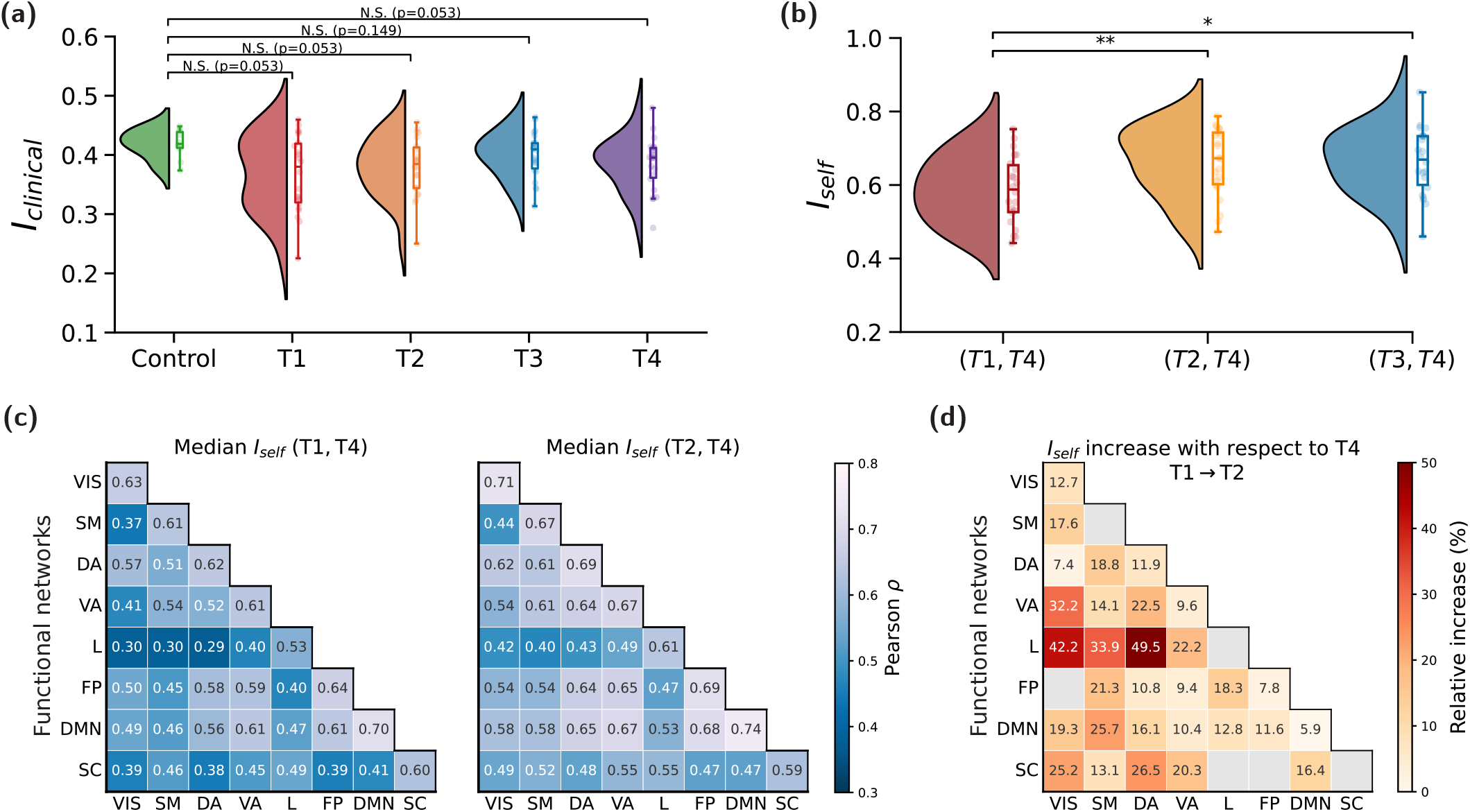
Longitudinal evolution of clinical and functional connectome identification in the complete-case sub-sample. Analyses replicate Fig. 2 using only patients with resting-state fMRI available at all four post-stroke sessions (*N* = 22), reducing statistical power but preserving the overall pattern of effects. **(a)** Clinical identifiability (*I*_clinical_), defined as similarity of each patient’s FC to the ECONS reference, remains reduced relative to the healthy control comparison (TrainStim vs. ECONS) across timepoints; in this smaller subsample, group differences are attenuated and some contrasts no longer reach significance, consistent with limited power. **(b)** Self-identifiability (*I*_self_ ) between earlier sessions and the chronic endpoint (T4) shows the same temporal profile as in the full cohort, with lower stability for the acute interval (*T*_1_–*T*_4_) relative to later intervals and a rapid increase by the early sub-acute stage. In this subsample, the early increase is weaker and does not consistently survive correction, but the direction of effect matches the main analysis (LMM with subject-specific random intercept; FDR-corrected). **(c)** Network-resolved *I*_self_ (median within/between Yeo networks and subcortical structures) reproduces the relative pattern of higher stability in association systems and lower stability in sensorimotor/subcortical blocks, albeit with increased variance. **(d)** Relative percentage increase in *I*_self_ from *T*_1_ to *T*_2_, referenced to *T*_4_, shows the same networks contributing most to early consolidation (notably VA, L, and SM), but with reduced statistical sensitivity compared to the full sample.

**FIG. S5.**
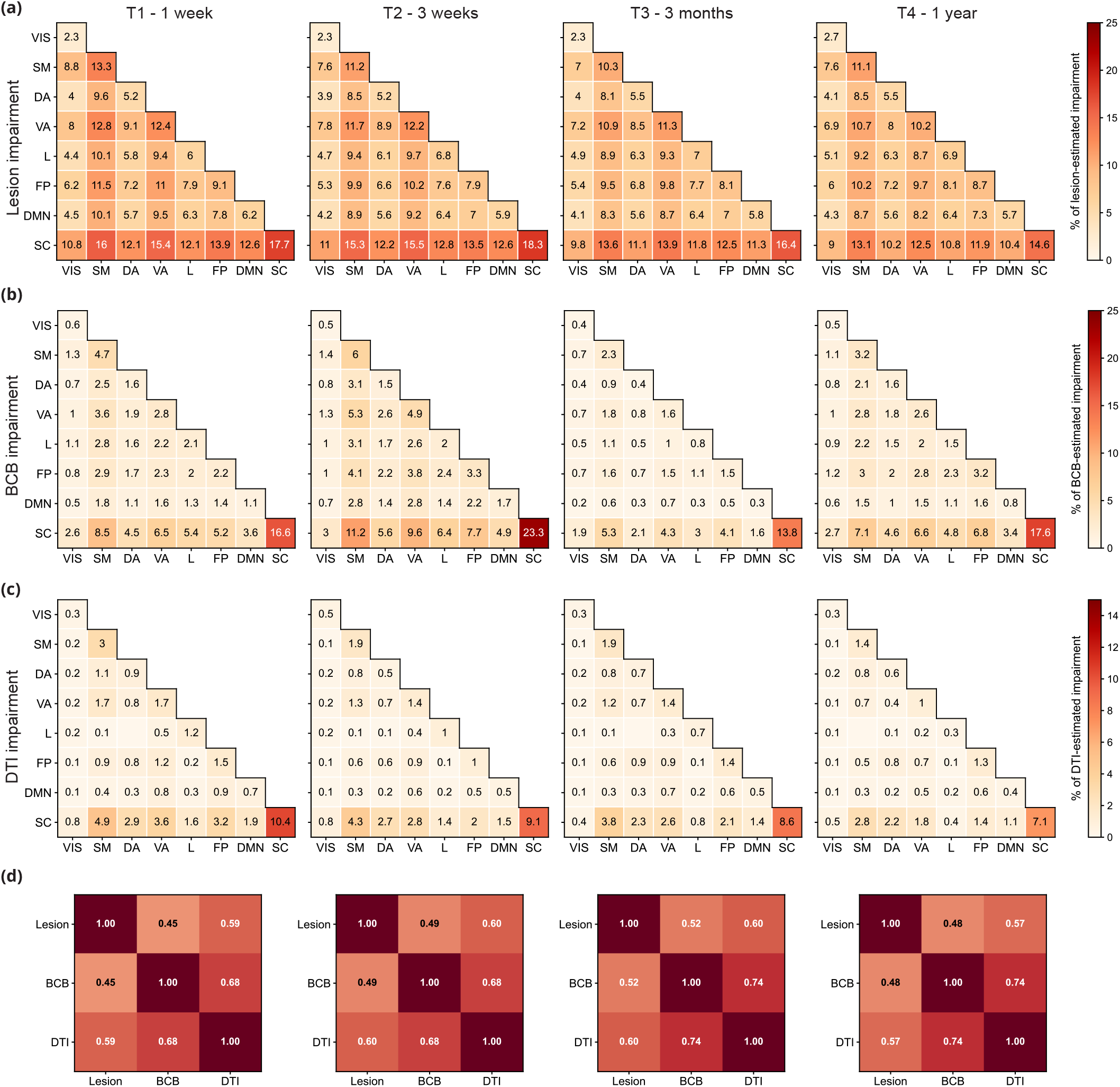
Comparison of direct and indirect measures of structural disconnection. Longitudinal quantification of structural network disruption for the same subset of patients used in Fig 3 estimated using three complementary methodological approaches across four time points (T1, 1 week; T2, 3 weeks; T3, 3 months; T4, 1 year). **(a)** Matrices display the network-level percentage of disconnected edges estimated via direct lesion masking (Lesion impairment), **(b)** probabilistic tractography using the BCB toolkit (BCB impairment), and **(c)** empirical patient-specific DTI tractography (DTI impairment). While the spatial topography of damage—heavily affecting subcortical (SC) and somatomotor (SM) connections—is consistent across methods, direct lesion masking yields higher absolute impairment magnitudes compared to DTI-based estimates. **(d)** Correlation matrices (Pearson’s r) quantifying the spatial similarity between the structural damage profiles generated by each method. High positive correlations confirm that the topological pattern of the disconnectome is robust to the specific modality used for estimation. Network abbreviations: VIS, Visual; SM, Somatomotor; DA, Dorsal Attention; VA, Ventral Attention; L, Limbic; FP, Fronto-Parietal; DMN, Default Mode Network; SC, Subcortical.

**FIG. S6.**
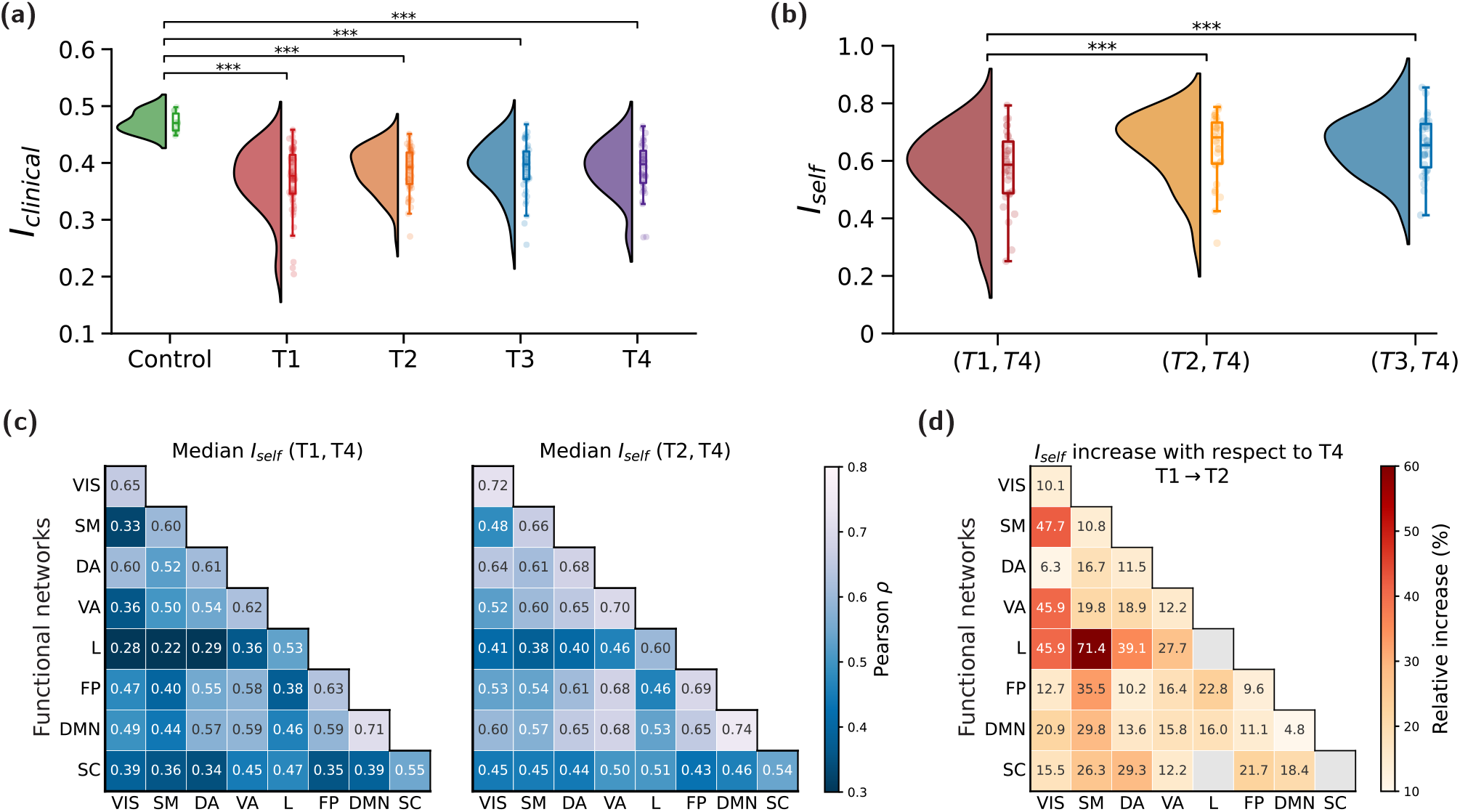
Robustness of clinical and functional connectome identification controlling for age, gender, and lesion size. Functional connectivity matrices were residualized for age and gender (all subjects) and log-transformed lesion volume (stroke patients only) prior to analysis. **(a)** Distribution of clinical identifiability scores (*I*_clinical_) for healthy controls (TrainStim vs. ECONS) and stroke patients (relative to ECONS). Controlling for covariates, stroke scores are significantly lower than healthy controls across all four time points ( ∗ ∗ ∗*p <* 0.001, FDR-corrected Mann-Whitney U test). **(b)** Distribution of self-identifiability scores (*I*_self_ ), quantifying patient connectome similarity between earlier time points and the chronic outcome (*T*_4_). Significantly lower *I*_self_ at *T*_1_ compared to *T*_2_ confirms that the rapid functional reorganization of the individual fingerprint within the first three weeks is robust to demographic and injury severity factors ( ∗ ∗ ∗*p <* 0.001, LMM). **(c)** Median *I*_self_ within and between seven canonical networks (VIS, visual; SM, somatomotor; DA, dorsal attention; VA, ventral attention; L, limbic; FP, frontoparietal; DMN, default mode) and subcortical structures (SC). **(d)** Significant relative percentage increase in *I*_self_ from acute to early sub-acute stages (*T*_1_ → *T*_2_), referenced to the chronic baseline (*T*_4_). Consistent with the main analysis, warmer colors denote networks undergoing rapid functional reconfiguration, notably the ventral attention (VA), limbic (L), and somatomotor (SM) systems (*p <* 0.05, LMM).

**FIG. S7.**
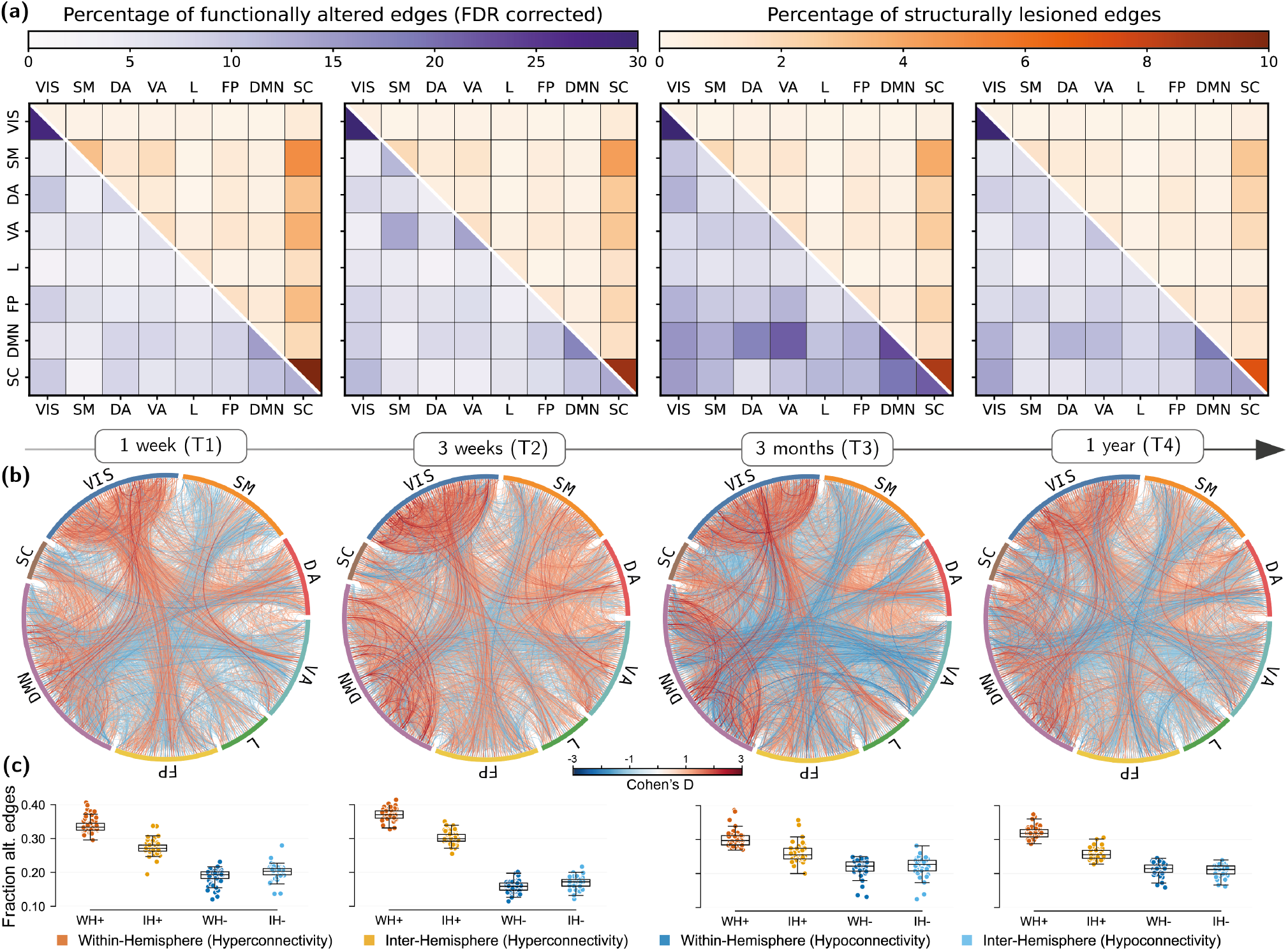
Robustness of longitudinal structural and functional alterations controlling for age, gender, and lesion size. Functional connectivity matrices were residualized for age and gender (all subjects) and log-transformed lesion volume (stroke patients only) prior to analysis. **(a)** Group-level matrices displaying the percentage of DTI-derived structurally lesioned edges (upper triangle) and covariate-residualized functionally altered edges (lower triangle; FDR-corrected Mann-Whitney U test, *q <* 0.05) relative to healthy controls across four time points (*T*_1_–*T*_4_). Consistent with the main findings, structural disconnection patterns remain static, whereas functional alterations evolve dynamically, confirming that the spatial dissociation between fixed structural injury and time-varying functional reorganization is independent of these covariates. **(b)** Circular plots depict positive (hyperconnectivity, red) and negative (hypoconnectivity, blue) functional alterations at *T*_1_–*T*_4_, quantified by Cohen’s *d* effect sizes from the residualized data. **(c)** Box plots summarize the fraction of altered edges classified as within-hemisphere (WH) or inter-hemispheric (IH), further subdivided into hyperconnectivity (*WH*+; *IH*+) and hypoconnectivity (*WH*− ; *IH*− ). Early stages (*T*_1_–*T*_2_) are dominated by within-hemisphere hyperconnectivity, whereas the late sub-acute/chronic phase (*T*_3_–*T*_4_) shows a relative increase in inter-hemispheric alterations, validating the shift from local to cross-hemispheric plasticity. Data are shown for the longitudinal subset (*n* = 22). Network abbreviations: VIS, visual; SM, somatomotor; DA, dorsal attention; VA, ventral attention; L, limbic; FP, frontoparietal; DMN, default mode; SC, subcortical.

**FIG. S8.**
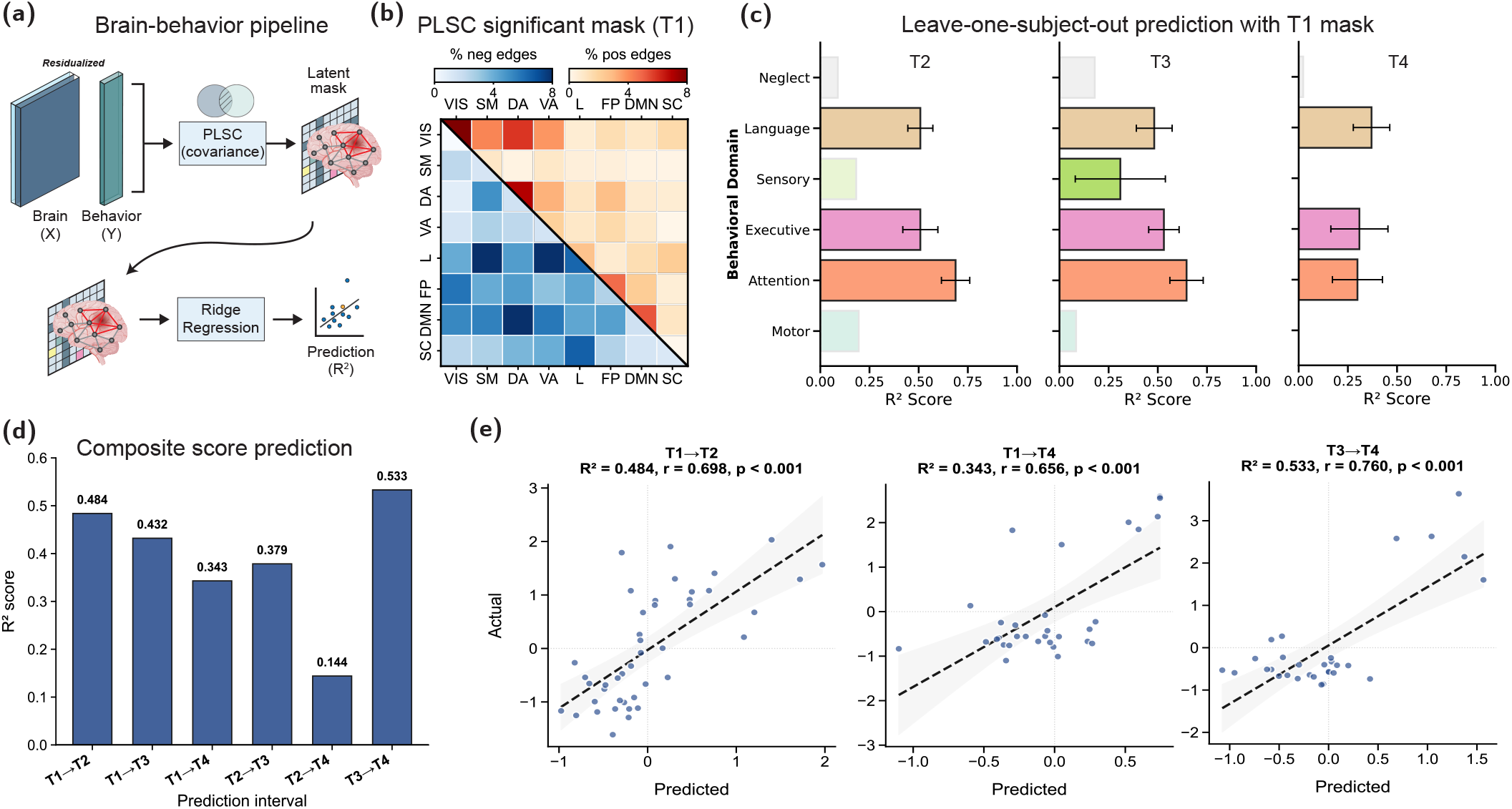
Robustness of multivariate brain–behavior mapping and longitudinal prediction controlling for age, gender, and lesion size. Functional connectivity matrices were residualized for age and gender (all subjects) and log-transformed lesion volume (stroke patients only) prior to analysis. **(a)** Two-stage predictive framework coupling Partial Least Squares Correlation (PLSC) with Ridge regression. Latent axes of maximal covariance between the residualized functional connectivity (Brain *X*) and behavioral scores (Behavior *Y* ) are identified via PLSC. Significant components generate a connectivity mask for Ridge regression, validated using a leave-one-subject-out (LOSO) approach to predict outcomes at subsequent time points (*T*_*i*_ → *T*_*i*+1…4_). **(b)** Network-level aggregation of the significant PLSC connectivity mask derived at the acute stage (*T*_1_) from the residualized data. Heatmaps display the percentage of significantly negative (blue) and positive (red) edges, highlighting the prominent contribution of higher-order association systems (FP, DMN) and subcortical structures (SC) to the acute behavioral phenotype. **(c)** LOSO prediction performance (*R*^2^) for individual behavioral domains using the *T*_1_ latent mask. Consistent with the main findings, the model successfully forecasts language, executive, and attention outcomes at follow-up (*T*_2_, *T*_3_, *T*_4_), whereas motor and neglect domains show negligible predictability from this functional signature. **(d)** Prediction accuracy for a global composite behavioral score across longitudinal intervals. Accuracy decays as the temporal gap from *T*_1_ increases but peaks during the late sub-acute transition (*T*_3_ → *T*_4_; *R*^2^ = 0.533), confirming that the late-stage stabilization of the brain–behavior relationship is robust to demographic and injury severity factors. **(e)** Scatter plots correlating actual versus predicted global scores for key intervals reported in panel (d) (e.g., *T*_1_ → *T*_2_, *R*^2^ = 0.484, *r* = 0.698, *p <* 0.001).

